# Fabrication, evolution, and mutual conversion of D-fucose-activatable and -repressible acetyltransferase upon mutations

**DOI:** 10.1101/2024.05.29.596397

**Authors:** Yuki Yanai, Miyu Tsukada, Yuki Kimura, Daisuke Umeno

## Abstract

The fusion of different proteins can result in the linkage-dependent emergence of molecular switches. In cases where allosteric regulation is designed between the input and output modules of fusion proteins, it is hard to predict whether on-switching or off-switching will occur. However, binding-induced folding, a non-allosteric molecular switch mechanism, has the potential to quickly establish a mutually regulatory relationship between the two fused proteins, in the way whether on-switching or off-switching will occur would be predictable. We inserted chloramphenicol acetyltransferase (CAT) from *E. coli* into a loop of a D-fucose-responsive mutant of transcription factor AraC, using linker libraries with various lengths. We found that on-switches tend to emerge when two proteins are fused with a small pitch gap at the junction, while fusion designs with a large pitch gap result in the frequent emergence of off-switches. Both types of switches rapidly evolved their switching efficiency upon mutations, establishing the D-fucose-on and -off regulation of CAT activity without disrupting the D-fucose-inducible logic of AraC function. To our surprise, both one-input/two-output split gates thus obtained could be easily inter-converted upon mutations. Through mutations, proteins not only frequently acquire properties as binding-induced folders, but also rapidly establish and evolve a mutual regulatory relationship with unrelated fusion partners, as well as transform their regulatory logic.

## Introduction

Different proteins are routinely fused to improve the rate of enzyme sequential reactions^1^, track spatiotemporal localization within cells and tissues^2^, stabilize expression^3^, and achieve protein purification^4^. With a linker with sufficient flexibility and length, proteins can be linked without losing their individual functions, yielding bifunctional proteins. On the other hand, linkage design can become more complex when establishing the mutual regulation of functions^5^.

In natural motifs such as transcription factors^6^, kinases^7^, and calmodulins^8^, the input and output domains are connected in a way that the binding events of one modulate the function of the other. The input and output modules within families can be easily interchanged to create new molecular switches^9–12^, suggesting that the transmission mechanism between the two domains is highly conserved among families. At the same time, there have been attempts to combine evolutionarily unrelated protein motifs to enable the regulation of one by the other. If the molecular binding domain undergoes a conformational change upon binding to its target, and if the output domain is structurally linked in a manner that renders it responsive to the conformational changes in the input domain, an allosteric regulatory mechanism can be established^13,14^. In their pioneering work, Doi et al. successfully inserted TEM1 β-lactamase into green fluorescent protein (GFP), allowing the detection of β-lactamase inhibitory protein (BLIP) binding to the TEM1 domain through changes in GFP fluorescence intensity^15^. In another study, bacterial SsrA peptide, known as a degradation tag, was embedded in the C-terminal helix of the naturally occurring photoswitch protein LOV2, creating a photo-switchable binder after four cycles of positive and negative selection^16^. Furthermore, the insertion of binding proteins into the Y145 site of GFP/YFP is known to result in the creation of biosensors, whereby binding events cause changes in the protonation state of the fluorophores ^17^.

When completely unrelated proteins are fused in a mutually regulating manner, the resulting switching behavior can be either on*-*switch or off*-*switch. When calmodulin was inserted at position 145 of GFP, the randomization of its linking residues resulted in the emergence of both calcium-on and calcium-off sensors^18^. The randomization of linker residues in LacI ligand-binding domain (LBD)-PurR DNA-binding domain (DBD) chimeras also produced mutants in which the DBD was positively or negatively regulated by β-galactosidase^19^. Through the replacement of the cytoplasmic signaling domain of the *E. coli* Tar chemoreceptor with that of EnvZ (a member of the same protein family), an aspartate-induced on*-*switch was obtained^12^. On the contrary, the replacement of the cytoplasmic signaling domain of cyanobacterial photoreceptor Cph1 with that of EnvZ resulted in a light-induced off*-*switch^20^. Furthermore, the Tar-EnvZ chimeric receptor was shown to fluctuate between on*-* and off*-*switch upon site-directed random mutagenesis at the linking residues^21^.

Although allosteric inter-regulatory relationships can be generated through fusion, it is difficult to predict whether the resulting fusion proteins will act as on*-* or off*-*switches. Furthermore, it is well-known that point mutations can often reverse the behavior of allosteric switches^22–24^.

Binding-induced folding (BIF) is an alternative principle for creating protein switches that bypass the need for designing conformational changes^25^. Here, by moderately destabilizing fused protein A so that it requires ligand-binding to properly fold, the function of its binding partner (protein B) also becomes conditional to the BIF of protein A. This way, various protein functions, such as those of Cas9 (Ref. ^26^), other enzymes^26^, fluorescent proteins^26^, and transcription factors^27,28^ have been demonstrated to be on*-*switched upon the binding of their ligand partners. Additionally, off*-*switches can be implemented by adopting mutually exclusive folding (MEF). Radley et al. attempted to directly insert Ubiquitin (Ub)^29^, whose N- and C-termini are at the distance of 38.5 Å, into a short loop of Barnase (Bn) from *Bacillus amyloliquefaciens*. Due to the incompatibility of the folding of both Bn and Ub, the ligand-induced folding of Bn upon its interaction with the protein Barstar triggered the unfolding of Ub, thereby establishing the Barstar-induced off*-*switching of Ub-folding. Thus, proteins with BIF properties can be harnessed for the on- or off-switching of other proteins.

In this study, we inserted chloramphenicol acetyltransferase (CAT) into the LBD of the D-fucose-responsive mutant of the L-arabinose-responsive transcription factor AraC (AraC_I46V_)^30^ (**Figure 1a**). The aim was to positively and negatively regulate the cellular function of CAT through D-fucose-induced AraC folding. We found that when the pitch gap at the fusion site was small, the fusion protein tended to become an on*-*switch, and when it was large, it tended to become an off*-*switch. Both types of D-fucose-induced CAT controls rapidly evolved by random mutagenesis, yielding bifunctional proteins with one-input two-output split gates. We found that even among populations having undergone multiple generations of evolution as D-fucose-on and -off switchers of CAT, variants with reverted behavior frequently emerged upon the introduction of random mutations (**Figures 1b and 1c**). Thus, random mutagenesis not only frequently confers proteins properties as BIF but could also rapidly establish mutually regulating relationships between two fused proteins, improving, or even reverting the properties of the generated molecular switches.

**Figure 1.**
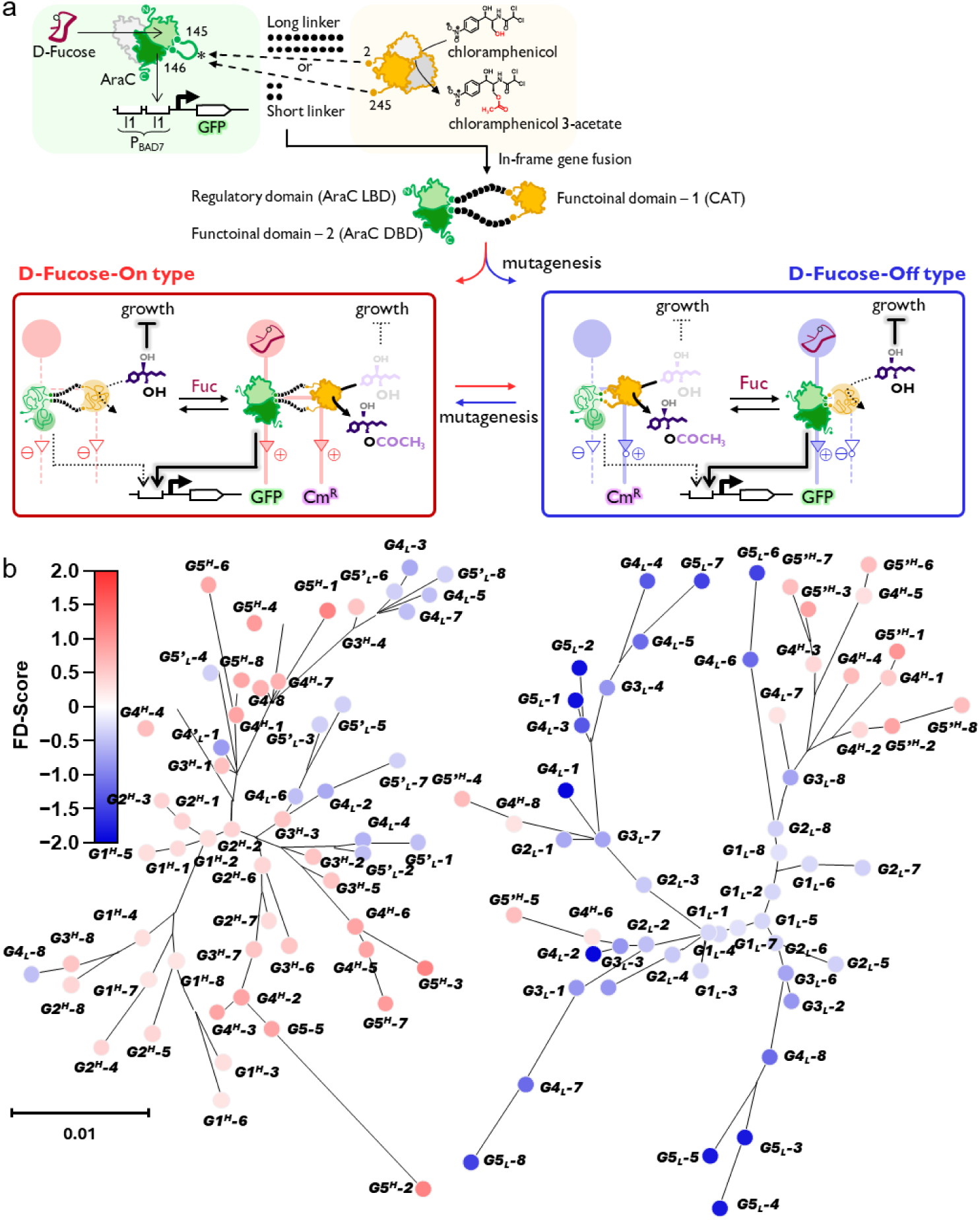
Development of D-fucose-regulated AraC::CAT fusion proteins and their interconversion upon mutagenesis. **(a)** One-input/two-output split gate proteins created by fusing chloramphenicol acetyltransferase (CAT) and sugar-responsive transcription factor AraC. In-frame fusion of CAT into the ligand-binding domain of AraC, followed by mutagenesis and screening for switching properties of CAT and for AraC. **(b)** Phylogenic analysis of D-fucose-on and -off switcher variants of AraC::CAT fabricated with long linkers with (9 + 9) spacers. The phylogenetic tree of the variants was created by MEGA^50^. In order to accommodate the space, the superscripts indicating Spacer size and the H/L symbols indicating high and low FD-score are omitted from the mutant name, and the name is simplified and the X-generation mutant Y is denoted as GX-Y. Primed G5 strands for the variants selected for reversed switching phenotype. From each generation, eight D-fucose dependency (FD) scores the mutant had. The variants with primed symbols represent mutants were acquired by screening for reversed phenotypes. Color of each variant represents its FD scores (for detail, see Fig.2). **(c)** Phylogenic analysis of D-fucose-on and -off switcher variants of AraC::CAT fabricated with (0 + 0) linkers. Scale bar refers to phylogenetic distance of 0.01 nucleotide substitutions per site.

## Results

### Evaluation of the switching properties of AraC::CAT fusion variants with different linkers

Chloramphenicol *O*-acetyltransferase (CAT) is the enzyme that acetylates the antibiotic chloramphenicol (Cm), thereby mitigating its inhibitory effects on bacterial translation machineries^31^. In this study, we attempted to insert the gene encoding this enzyme (*cat*), in-frame, into the LBD of *araC*^32^ derived from *E. coli*, to establish sugar-inducible CAT.

As the insertion point for CAT, we chose residue 145 in the small loop region located on the surface of the LBD of AraC. Being located on the opposite side of the dimer-forming surface of AraC, it is sufficiently distant from both ligand-binding and DNA-binding moieties. Molecular models created by MODELLER^33^ and/or AlphaFold2^34^ indicated that CAT can be fused without geometrically blocking CAT trimer formation or DNA-binding by AraC (**Figure 1A**). To monitor AraC function, we adopted the P_BAD7_ variant of the L-arabinose-induced promoter, which we recently found to be the ideal reporter of the BIF of AraC^35^. We also introduced an isoleucine-to-valine mutation in residue I46 of AraC, which increased its binding affinity to D-fucose, originally known as the antagonist of AraC^36^.

In the structural model, the distance between the alpha carbon atoms of residues 145 and 146 in AraC was 3.9 Å, whereas the distance between the alpha carbons of residues 2 and 219 was 24.1 Å. To accommodate the large pitch gap between the two at the fusion point, we introduced a relatively long and flexible linker consisting of nine residues at both CAT insertion points. This prototype AraC::CAT exhibited the functions of both AraC (D-fucose-induced activation of P_BAD7_-GFP) and CAT (ensuring the cell growth of host cells in the presence of Cm) (**Figure S1**). Cells harboring this prototype AraC::CAT displayed slightly higher Cm-resistance in the presence of D-fucose and L-arabinose, implying that CAT is positively regulated by the binding of D-fucose to AraC. Two codons neighboring *cat* were randomized both at the linkers, yielding a “(9 + 9) linker library” with 32^4^ ∼ 1 × 10^6^ theoretical diversity (**Figure 2a**). In search for the optimal linker length for acquiring D-fucose-on/off switchers, we incrementally truncated the AraC sides of the (9 + 9) library, preparing 3 additional libraries (the (6 + 6), (3 + 3), and (0 + 0) libraries). In addition, four residues on the CAT side of the (0 + 0) linker were removed to produce the “(−4 + −4) linker library.”

**Figure 2.**
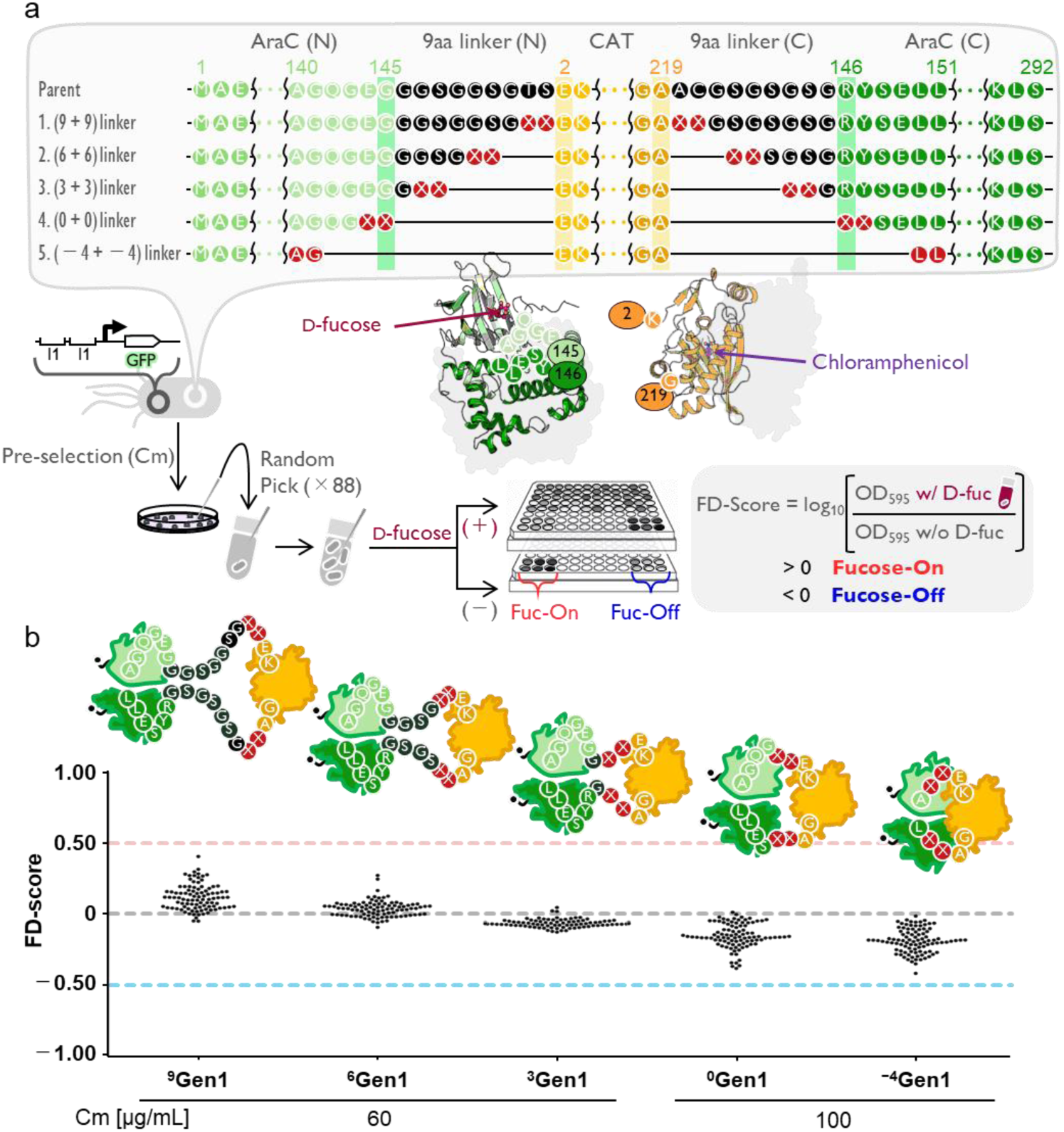
Distribution of the D-fucose dependency of the CAT activity, which is inserted into the ligand-binding domain of AraC using linker libraries of different lengths. (a) Workflow for creating and evaluating the AraC::CAT variant libraries. *E. coli* JW0063 cells harboring the P_BAD7_-GFP plasmid were transformed with each of the five indicated linker libraries. From the variants that retained detectable CAT activity either in the presence or absence of D-fucose, 88 clones were randomly picked and grown on LB-Cm liquid medium with or without D-fucose. The ratio of cell density (OD_595_) in the presence/absence of D-fucose was calculated for each variant and violin-plotted to obtain their respective FD scores. See the **Methods section** for further details. (b) Distribution of FD scores of AraC::CAT (145) variants of linker libraries. The common logarithm of the ratio of OD_595_ of randomly picked 88 transformants were determined. The Cm concentration in the medium was 60 µg/mL for the (9 + 9), (6 + 6), and (3 + 3) linker libraries and 100 µg/mL for the (0 + 0) and (−4 + −4) linker libraries.

Each of the AraC::CAT libraries was independently introduced into *E. coli*, and the clones that could endow Cm-resistance were selected on plates supplemented with D-fucose. We found that a significant fraction of all five libraries lost CAT activity (**Table S1**). Therefore, these libraries were pre-selected using Cm to remove dead mutants. In the case of the (9 + 9) linker library, 88 CAT-positive variants were randomly picked from the D-fucose (+)/Cm (+) plate, followed by the evaluation of the D-fucose dependence of their CAT activity. To compare the switching behaviors of AraC::CAT variants, we defined the D-fucose dependency score (FD score) as the common logarithm of the ratio of cell density with/without D-fucose. Most variants in the library exhibited FD scores higher than 0, indicating that their CAT activity increased upon D-fucose addition. With this linker length, the folds of AraC and CAT are compatible with each other, so that the stabilization of AraC by D-fucose binding also has a positive effect on the fusion partner (CAT). We sequenced eight (9 + 9) linker variants with high FD scores but found no indication of any enriched sequences (**Tables S2 and S3**).

As the length of the linker decreases, the distribution of the FD scores gradually decreased. Among variants of the (3 + 3), (0 + 0), and (−4 + −4) libraries, the majority exhibited negative FD scores (**Figure 2b**), indicating that the folding of AraC and CAT is not compatible with these linkers and that the stabilization of AraC by D-fucose binding negatively impacts CAT folding stability. Here again, all the eight variants with the lowest FD scores were unique in sequence (**Table S4**). We could not extract any apparent sequence characteristics discriminating switchers from non*-*switching variants with FD scores near zero (**Table S5**).

### Directed evolution of the D-fucose*-*on switcher variant

In general, most amino acid mutations introduced into proteins decrease structural stability, with a wide range of destabilization degrees^37^. This destabilizing effect of mutations converts proteins into BIFs, sometimes with surprisingly high frequency^38^, rooted in the ubiquitous fact that the gain of free energy from interactions contributes to the stabilization of protein structure. With this mode of action as a working principle, various binding proteins have been successfully used as sensory components^25–28^.

Considering the destabilization effect of protein insertion into the structure of AraC, it is suggested that the D-fucose-induced increase in CAT activity, which was exhibited by many members of the (9 + 9) linker library, is a read-out of the BIF of its binding partner AraC through interactions with D-fucose. We hypothesized that if this is the case, a further reduction in the folding stability of AraC through random mutagenesis would further increase the degree of its D-fucose dependency, thereby improving the switching efficiency of CAT activity.

Eight variants exhibiting the highest FD scores (0.2377 to 0.4077) among the 88 randomly picked variants from the (9 + 9) linker library was selected as ^9^Gen1_H_-variants, mixed, and subjected to error-prone PCR (epPCR) to randomly introduce amino acid substitutions into the fusion protein coding region. The resultant library was transformed into *E. coli* and the transformant clones that could grow on Cm-containing agar plates were pre-selected in the presence of D-fucose (10 mM) to remove the variants without CAT activity. From the clones that could form colonies on agar containing 60 µg/ml of Cm, 88 variants randomly selected (^9^G2) were to evaluate the D-fucose dependency of their CAT activity. The eight variants with the highest FD scores were identified as ^9^Gen2_H_ and subjected again to whole gene epPCR and screening for high FD scores (^9^Gen3_H_). By selecting as many as eight variants from a population of only 88 (with 9% being the parents of the next generation), we minimized the possibility of acquiring and concentrating rare mutations that confer high switching performance by evolving a tried-and-true allosteric mechanism. In this scheme, the evolution of the FD score, if it takes place, is most likely to occur only through high-frequency mutations, including destabilization mutations that could make AraC::CAT folding dependent on the D-fucose-induced stabilization.

The sequencing of the selected variants from each generation (**Figures S2–S4**) revealed the progressive accumulation of multiple amino acid substitutions over generations. Most strikingly, many of these mutations were accumulated primarily in the LBD of AraC (**Figure 3b**). When calculated using FoldX^39^, most mutations were predicted to have destabilizing effects on AraC (**Figure S5**). These mutations are likely to increase the degree of dependency of AraC folding on the stabilization achieved through its interaction with D-fucose.

**Figure. 3.**
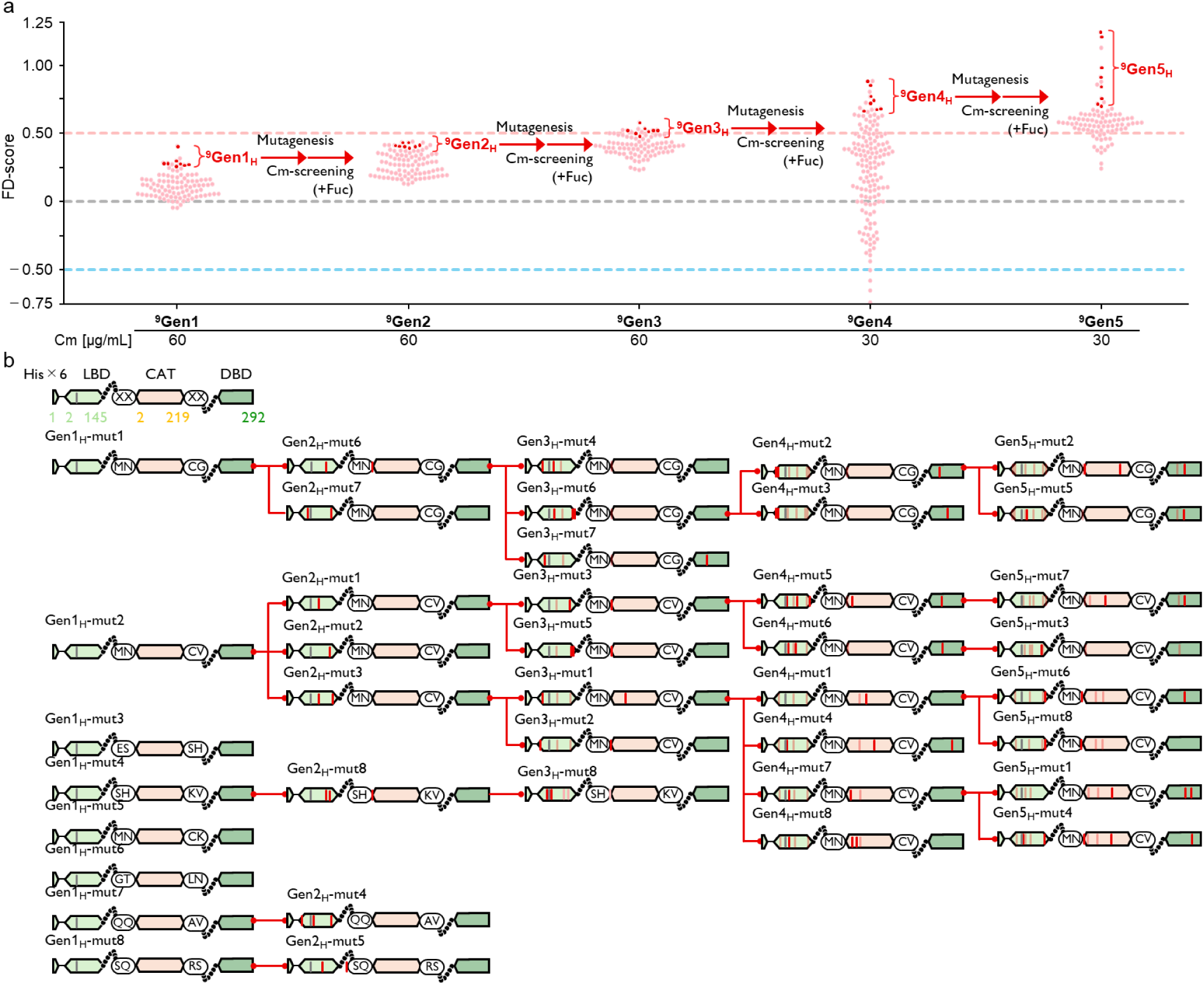
Directed evolution of D-fucose-ON switching properties of AraC::CAT variants with 9+9 linkers. (a) Transition of FD score distribution of AraC::CAT variants with (9 + 9) linkers over generations. Eight selected variants with the highest FD scores (indicated as red lines) were selected and pooled as parents for the next round of mutagenesis and screening. (b) Transition of genotypes of the selected variants in each generation. Red arrows indicate the evolutionary paths, and newly introduced amino acid mutations are indicated as red vertical lines. The two circled letters represent the amino acid sequences of the randomized residues in the initial linker library (^9^Gen1). For the detailed sequences, see **Figures S2-S4, S6, and S7**.

We further conducted a 3^rd^ round of random mutagenesis and screening but could not find any variants with improved FD scores, at least in the 88 variants tested. Thus, in the fourth generation, we reduced the selection pressure for CAT activity by decreasing the concentration of Cm (30 µg/ml instead of 60 µg/ml). We expected that slightly lower selection pressure would allow the emergence of mutations that would further decrease the stability of CAT and increase its functional dependence on the folding integrity of AraC. Indeed, we could observe further improvements in FD scores, enabling two additional rounds of whole-gene mutagenesis and screening. The highest FD score achieved was 1.246 (17.6-fold D-fucose-induced increase in growth in the presence of Cm) (**Figure 3a**).

The resultant high FD scoring mutants (**Figures S6 and 7**) displayed highly destabilizing mutations in the CAT region, as the calculated *ΔΔG* values ranged from +10.8 to +17.1 kJ/mol. These are similar to the expected stabilization effect of D-fucose binding (12–17 kJ/mol in *Δ*G_bind_), as assumed from the reported affinity between D-fucose and wild-type AraC (∼5 mM)^40^.

### Directed evolution of D-fucose*-*off variants

When a guest protein is inserted into the loop of a host protein and the host and guest proteins are joined without linkers that are necessary for filling the pitch gap between the two, their folding becomes incompatible. In this case, the structural stabilization of one component compromises the folding of the other^29,41,42^. This is consistent with the fact that the distribution of FD scores of AraC::CAT variants is reduced when adopting short linkers (**Figure 2b**). Most variants in the 0 aa linker library exhibited negative FD scores, and we believe they likely behaves as MEF. If this was the case, we hypothesized that the off*-*switching efficiency of CAT activity could be quickly improved through the accumulation of destabilizing mutations, which should enhance the BIF of the input module (AraC).

We selected eight mutants with the lowest FD scores (−0.3781 to −0.2772) from 88 AraC::CAT variants in the (0 + 0) linker library (**Figure 2b**). The plasmids encoding these eight variants (^0^Gen1_L_) were mixed and subjected to whole-gene random mutagenesis using epPCR. *E. coli* transformed with the resultant AraC::CAT library were grown in a D-fucose-free medium containing Cm. From the clones that could form colonies on the plates, 88 variants were randomly selected and subjected to screening for low FD scores. The eight with the lowest FD scores (^0^Gen2_L_: −0.5784 to −0.4311 in FD scores) were selected and pooled as the parents for another round of mutagenesis and screening for lower FD scores. This mutagenesis and selection cycle was repeated for three more generations, yielding ^0^Gen3, ^0^Gen4, and ^0^Gen5 variants (**Figure 4a**).

**Figure 4.**
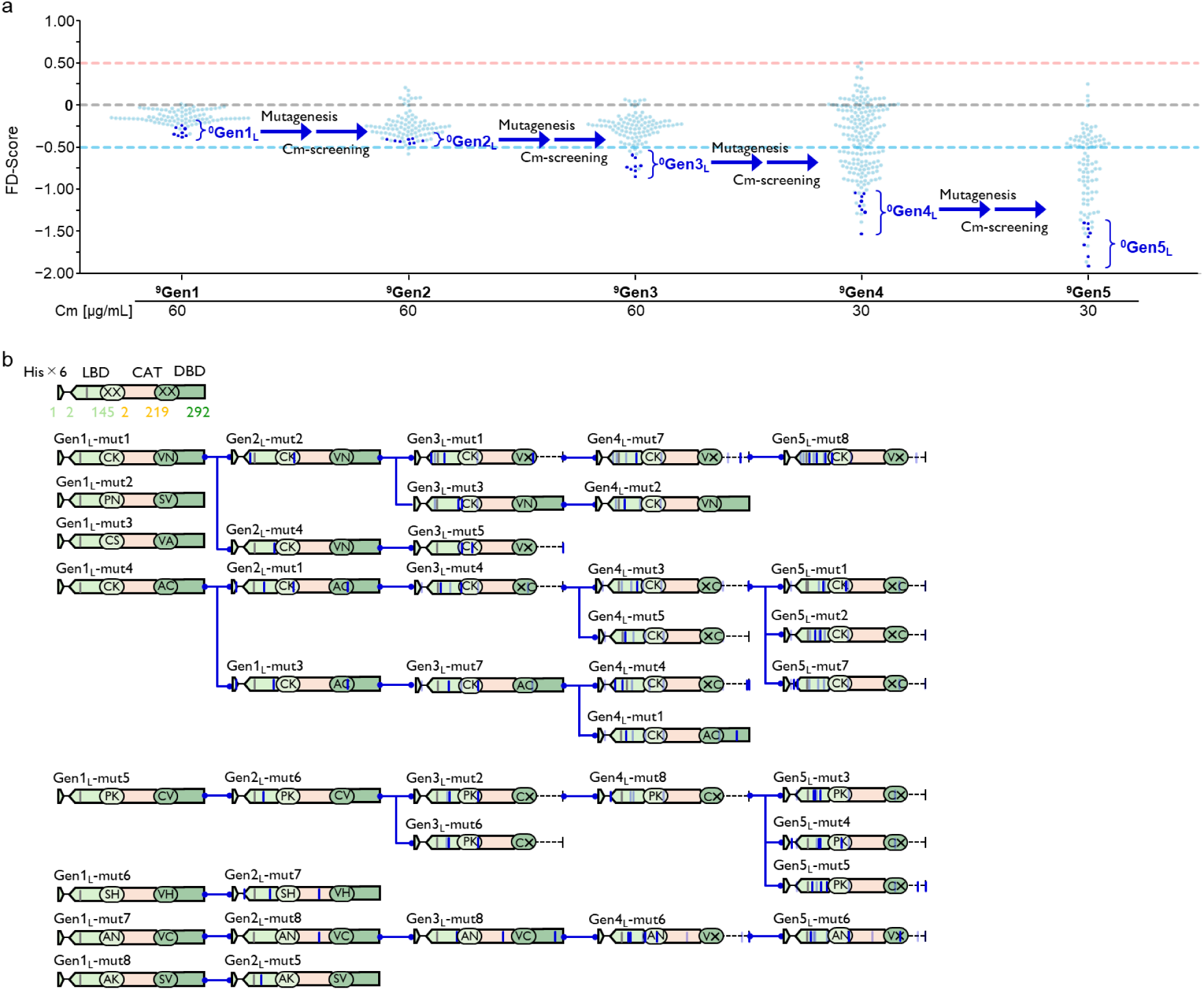
Directed evolution of D-fucose-off switching properties of AraC::CAT variants with (0+0) linkers. (a) Transition of the FD score distribution of AraC::CAT variants with (0 + 0) linkers over generations. Eight selected variants with the lowest FD scores (indicated as blue lines) were selected and pooled as parents for the next round of mutagenesis and screening. (b) Transition of genotypes of the selected D-fucose-off variants in each generation. Blue arrows indicate the putative evolutionary paths, and newly introduced amino acid mutations are indicated as blue vertical lines. The two circled letters represent the amino acid assigned at the randomized residues in the initial (0+0) linkers (see Figure 2A). For detailed sequences, see **Figures S10–S14.**

In the fourth round of directed evolution, we observed a marked decrease in the fraction of variants that could form colonies on agar plates containing 100 µg/ml Cm (**Table S1**). For rapidly evolving populations of proteins, there is a known need for acquiring stabilizing mutations that compensate for the destabilizing effect of the continuous accumulation of mutations^43^. This phenomenon is likely to be prevalent during the evolution of switching properties based on BIF, as it pursues a decrease in protein folding stability. To address this “mutational dead end,” we opted to decrease the selection pressure on CAT activity, simply by decreasing the concentration of Cm added to the agar plate. By adopting the Cm concentration of 60 µg/mL instead of 100 µg/mL, the distribution of FD scores started to progressively decrease again with each generation. By the fifth generation, mutants with over 100-fold higher signal-to-noise ratios were obtained (**Figure 4a**).

Throughout the evolution of low FD scores, we encountered the frequent emergence of truncated mutants. In the 2^nd^ generation, three variants exhibiting particularly low FD scores were truncated by acquiring nonsense mutations downstream of *cat* (at positions 146 and 147 in AraC). Due to the loss of ∼50% of the AraC sequence, including the entire DNA binding domain (DBD), all these mutants completely lost function of AraC (sugar-induced transcription activation of P_BAD7_) (**Figure S8**). To access eight off*-*switching mutants that retained the entire frames (designated ^0^Gen2_L_), we had to sequence 17 mutants with low FD scores. Despite starting from non-truncated parents (^0^Gen2_L_), five truncation mutants were found among the eight tested mutants with the lowest FD scores in ^0^Gen3. This frequent occurrence of truncation is likely due to the scoring system we adopted for defining better off*-*switchers (i.e. low FD-score). It is likely that the most “accessible” solution to achieve a higher FD score was to recover the on*-*signal value by removing 51% of the C-terminal part of AraC, likely by increasing the degrees of freedom (**Figure S9**). CAT is known to form reaction pockets at the homotrimer interface for acetyl transfer activity^47^. The restoration of the conformational freedom of the C-terminal region may have improved the ease of assembly of CAT trimers. Most importantly, however, about a third of AraC::CAT variants achieved considerably low FD scores (off*-*switching properties) without losing the entire reading frame, yielding bifunctional variants with both CAT and AraC functions.

Regardless of whether the entire frame was retained, mutations found in good switchers (those with the lowest FD scores) were considerably similar (**Figures S10–S14**). We observed the accumulation of very few mutations in the reading frame of CAT and the DBD of AraC. Most of the accumulated mutations were identified in the AraC LBD. FoldX calculations confirmed that most of the destabilization (*ΔΔG* ranging from +19 to +20 kJ/mol for eight ^0^Gen5_L_ variants) of AraC::CAT arises from mutations in AraC’s LBD. This suggests that D-fucose-off variants also mainly operate through BIF. The tendency for a biased accumulation of mutations in the AraC LBD region (**Figures S10–S14**) was even more pronounced for D-fucose-off switches than for the D-fucose-on switches (**Figures S2-S4, S6, and S7**), likely due to the stronger selection pressure to maintain CAT integrity.

### Dual functionality of the D-fucose-on and -off variants

Many of the variants in the D-fucose-on lineage (**Figure 3**) retained AraC’s function as a transcription factor, despite the absence of selection for preserving its function and the presence of significant selection pressure on the stability loss of the LBD. As a result, we acquired dual-functional AraC::CAT variants whose two functions responded positively to D-fucose (^9^Gen5_H_-mut1 in **Figure 5**). This mutant activated P_BAD7_ in response to D-fucose (**Figure 5a**), and allowed host cells to grow in a D-fucose-dependent manner. With four mutations accumulated on the LBD (calculated to achieve a destabilization of 17 kJ/mol, (**Figure S15**), considering that the P_BAD7_ promoter was designed to respond to the effective concentration of AraC rather than to its structural change (allostery) upon ligand binding^30,35^, AraC’s LBD is likely to become a binding-induced folder, and its D-fucose-induced folding could trigger folding of its fusion partners, AraC’s DBD and CAT, with four mutations accumulated on the LBD (calculated to achieve a destabilization of 17 kJ/mol, (**Figure S15**)).

**Figure 5.**
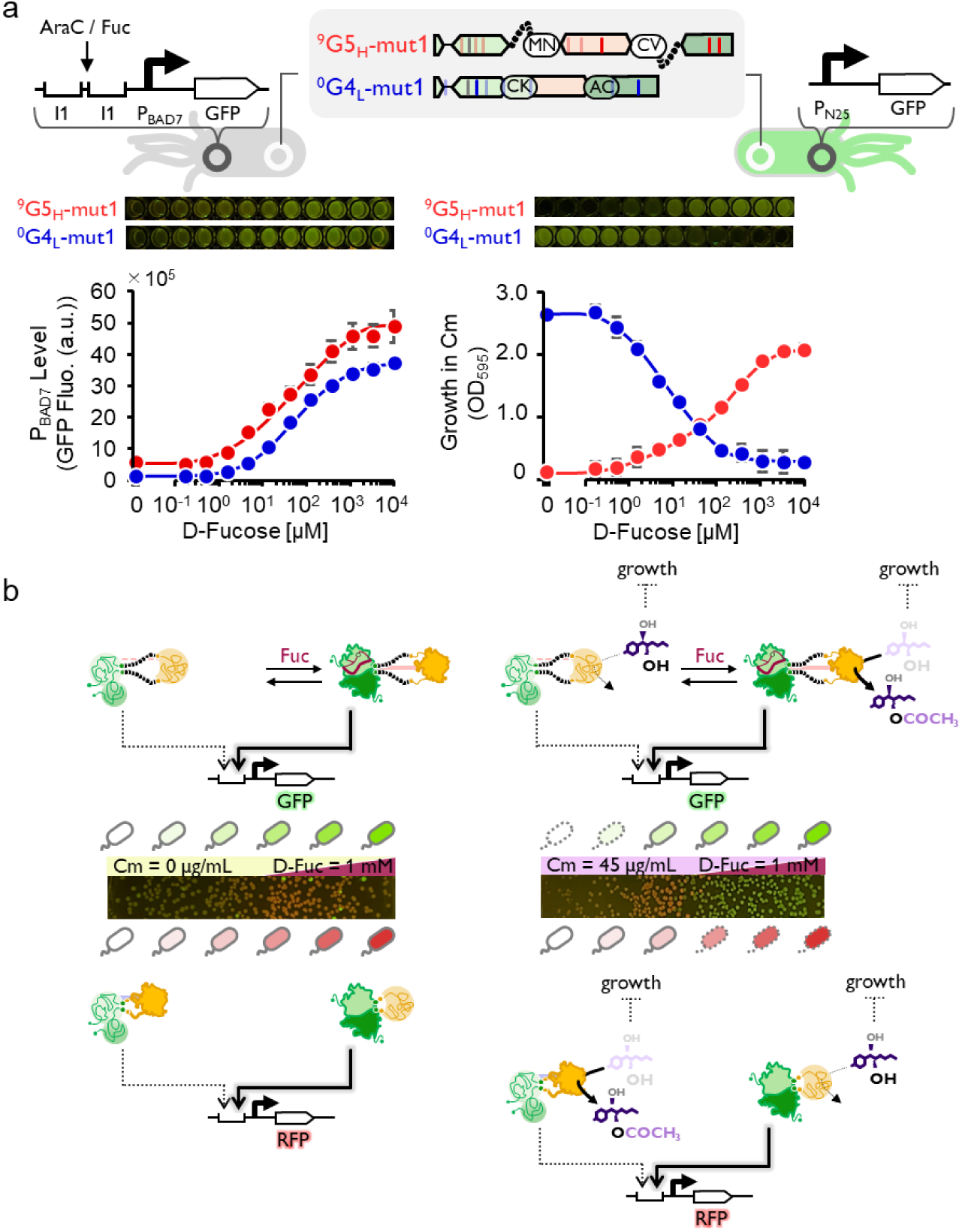
Simultaneous and independent control of AraC::CAT functions. (a) Transfer function of AraC function (activating efficiency of P_BAD7_ promoter) to D-fucose in representative variants (^9^Gen5_H_-mut1 and ^0^Gen5_L_-mut1) of D-fucose-on and - off evolutionary lineages. The data corresponds to the mean ± s.d. of four biological replicates. (b) Transfer function of CAT activity (growth on Cm-containing medium) to D-fucose in representative variants (^9^Gen5_H_-mut1 and ^0^Gen5_L_-mut1) of D-fucose-on and -off lineages. The data corresponds to the mean ± s.d. of four biological replicates. (c) Morphogenic control of the macroscopic pattern of the mixed cells transformed with the D-fucose-on (harboring P_BAD7_-GFP) or D-fucose-off (harboring P_BAD7_-mCherry) variants in response to the D-fucose concentration gradient.

The evolution of the D-fucose-off phenotype (lower FD scores) (**Figure 4**) led to the accumulation of nonsense mutations downstream of the CAT of some variants, which lost its function as a transcription factor (**Figure S8**), while many of the AraC::CAT retaining the whole frame could activate transcription of P_BAD7_. These mutants activated the transcription of P_BAD7_-GFP in response to D-fucose, as shown in (**Figure 5a)**, whereas their CAT function was inversely regulated (D-fucose-repressed, **Figure 5b**). The ^0^Gen4_L_-mut1 exhibited higher stringency in regulating the P_BAD7_ promoter expression than the ^9^Gen5_H_-mut1 (lower P_BAD7_ induction in the absence of D-fucose, **Figure 5a**), suggesting that its more dependent on the enthalpy gain upon D-fucose-binding to AraC_LBD_. Likely due to pronounced (laboratory-evolved) incompatibility at the junction region over generations, the D-fucose-induced folding of AraC_LBD_ compromises the functional structure of its fusion partner CAT, leading to a dose-dependent reduction in its activity.

Each of these mutants can be considered a “protein circuit” with a distinct logic. When cells expressing these mutants are initially plated in a mixed state, the colony-forming spaces become segregated by creating a D-fucose concentration gradient. This results in the formation of characteristic macroscopic patterns corresponding to the D-fucose concentration gradient (**Figure 5c**).

### Evolution of the reverse switching behaviors in the D-fucose-on/off lineage

AraC::CAT variant population members with long (9 + 9) linkers were initially evolved for D-fucose-on behavior (high FD score), and all members exhibited D-fucose-on phenotypes in ^9^Gen2 and ^9^Gen3 populations. Likely due to the destabilization caused by accumulated mutations, we had to lower the concentration of Cm to continue improving the FD score at ^9^Gen4. Interestingly, we found that a minor fraction of ^9^Gen4 populations exhibit D-fucose-off phenotypes (^9^Gen4_L_ variants in **Figure 3a**). To test whether this reverted phenotype could be improved by mutations, we picked eight variants with the lowest FD scores from the 88 variants of the fourth generation of the D-fucose-on lineage (^9^Gen4_L_ in **Figure 6a**) and subjected them to whole-gene mutagenesis by epPCR. From the resultant library, which was pre-selected for Cm-resistance in the absence of D-fucose, 88 variants (^9^Gen5’) were randomly picked, and their FD scores were analyzed. Although the proportion of D-fucose-off mutants increased, we could not obtain a mutant with a lower FD score than its parent, ^9^Gen4_L_. The switching property of a D-fucose-on mutant in the (9 + 9) linker lineage (**Figure 3**) is likely to have evolved by devaluing the CAT function to zero in a ligand-free environment **(Figure S16**). Many mutations would have already accumulated in the CAT gene at the ^9^Gen4/^9^Gen5 stages (**Figure S6** and **S7**). Therefore, it was surprising that the D-fucose-off phenotype also emerged in this line. If the stability of these fusion proteins have already been well down-tuned, the function as a D-fucose-off switch that appears here would not be possible without regaining the intactness of CAT in the ligand-free state.

**Figure 6.**
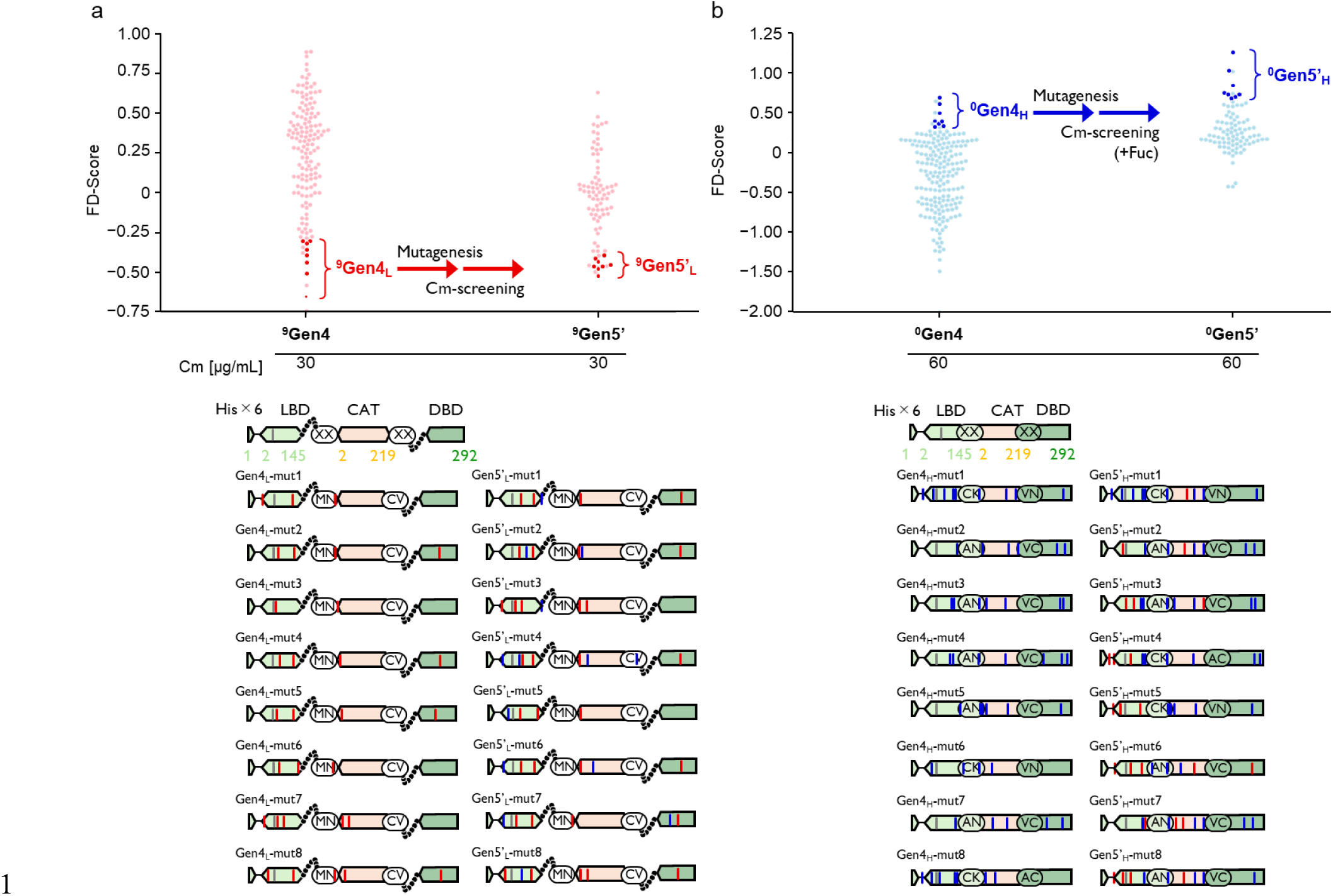
Emergence of variants with reverted switching properties from (A) D-fucose-on and (B) D-fucose-off parents upon random mutagenesis. (a)From the 88 members of AraC::CAT variants with (9 + 9) linkers (^9^Gen4, D-fucose-on lineage), eight variants with the lowest FD scores (^9^Gen4_L_) were pooled, mutagenized, and pre-selected for the retention of CAT activity in the absence of D-fucose. 88 variants (^9^Gen5’) were randomly selected and evaluated for their FD scores, among which eight variants with the lowest FD scores were named ^9^Gen5’_L_. (b) Eight variants with the highest FD scores (^0^Gen4_H_) were pooled, mutagenized, and pre-selected for the retention of CAT activity in the presence of D-fucose. From the 88 members of AraC::CAT variants with (0 + 0) linkers (^0^Gen5’, D-fucose-on lineage), 88 variants (^0^Gen5’ variants) were randomly selected and evaluated for their FD scores. Eight variants with the highest FD scores among them were named ^0^Gen5’_L_.

All through the course of evolving off-switchers, in the lineage of AraC::CAT variants with short (0 + 0) linkers (**Figure 4A**), the D-fucose-on variants (with reverted phenotype) emerged. From the 88 variants of the fourth generation of the D-fucose-on lineage, eight variants with the highest FD scores (^0^Gen4_H_) were selected (**Figure 6b**), pooled, and subjected to epPCR to introduce random mutations. After pre-selection for Cm-resistance in the presence of D-fucose, 88 variants were randomly picked and tested for FD scores. This time, we obtained numbers of variants with evidently increased FD scores (^0^Gen5’_H_). The highest FD score among the 88 mutants tested was 1.035 (^0^Gen5’_H_-mut1), which is comparable to the score shown by the highest D-fucose-on mutant ^9^Gen5_H_-mut1, acquired after four rounds of evolution. (**Figure 3**).

## Discussion

In the case of natural transcription factors including AraC and LuxR, LBD and DBD have evolved tight connections, whereby ligand-binding events in the LBD are transmitted to the DBD mainly through sophisticated allosteric mechanisms. However, even if these are maintained, their cellular behavior can be easily attenuated depending on the composition of the target sequence and/or the effective concentration of itself. For instance, when the AraC operator configuration becomes nonoptimal^30^ or when the stringency of LuxR is reduced by mutation^38^, these transcription factors start to act as “always-on”: their target genes are transcribed continuously, regardless of ligand molecule concentrations. However, through random mutations, their switching property can be quickly re-established. We have recently reported that random mutagenesis converts non-switching variants of the transcription activator AraC^30^ and LuxR^38^ into variants with excellent signal-to-noise ratios, sometimes at the frequency of 20%. Here, we showed that this evolution occurred through the destabilization of their folding, thereby conferring them BIF properties. The folding of the LBD and DBD is tightly coupled by a physical contact interface strengthened by evolution, and the ligand-induced folding of the LBD becomes a prerequisite to the adoption of a functional structure by its folding partner, DBD.

BIF^25–28^ has been proposed as an attractive approach for the rapid development of molecular switches and sensors without the time-consuming task of allosteric regulation design. Although there is no detailed information on the frequency of its occurrence, the establishment of a relationship in which molecular recognition (stabilization) of one protein triggers the adoption of a functional structure by the other can also occur in fusions of evolutionary unrelated proteins ^26–28^. We inserted CAT into AraC with various linkers and tested the degree of dependency of CAT function on AraC ligand-binding (**Figure 2**). With long linkers, CAT evolved D-fucose-dependent properties by accumulating mutations throughout whole protein sequence (**Figure 3**). Considering that the mutants obtained after 5 generations have 6–13 mutations (**Fig. S15:** calculated *ΔΔG* ranges from 28.65-29.66 kJ/mol), all these D-fucose-on mutants are believed to have an overall lowered stability, and the folding of CAT is seemingly dependent on the D-fucose-induced stabilization of AraC’s LBD.

When AraC and CAT are fused without a linker, the pitches at the junction of the two in the folding state do not match, so their folding is no longer compatible. As a result, they are supposed to exert negative effects on each other’s functions. Indeed, variants with low FD scores (D-fucose-off type) dominated the population in the (0 + 0) linker library (**Figure 2b**), and its signal-to-noise ratio evolved quickly (**Figure 4**). Here again, the LBD of the mutants is thought to have improved its properties as a binding-induced-folder during evolution. Amino acid mutations accumulate intensively in the LBD with each generation (**Figures S10–S14**). By the fifth generation, variants accumulated an average of 8.5 mutations (**Figure S14**), and their calculated stability effects reached up to 36.54–39.19 kJ/mol (**Fig. S15**). Additionally, the mutant ^0^Gen4_L_-mut1, which was the lowest FD scorer obtained in this evolutionary lineage, showed extremely low leak of AraC function in an environment without D-fucose, and as a result AraC was stringently behaved as D-fucose-on switcher (**Figure 5a**). In contrast, the CAT domain acquired only a few mutations throughout the initial generations. This is probably due to a strong selection pressure for maintaining CAT activity under ligand-free conditions, i.e., maintaining CAT integrity, as a prerequisite for the evolution of low FD scores. In contrast, many mutations accumulated in CAT in the later stages of the evolution of D-fucose-on switches in the long (9 + 9) linker lineage (**Figures S6 and S7**), as well as in the process of evolving reverted switchers from the zero-linker lineage originally bred for their D-fucose-off properties (**Figures S7, S17** and **S18**).

We found that BIF-based protein switches can be easily reversed simply by whole gene mutagenesis, even for populations that had been already well differentiated as D-fucose-on and -off switchers. From the population that evolved as D-fucose-on mutants for four generations, up to 26% of the variant pool exhibited D-fucose-off phenotypes when parented by low FD scorers of ^9^Gen4. Similarly, from the population subjected to selection for D-fucose-off behavior for 4 rounds, we identified 38% of D-fucose-on variants in the library prepared by mutagenizing high-scorers of ^0^Gen4 mutants. The neutral networks of the D-fucose-on and -off switchers are extremely intertwined, and there seem to be countless communication paths for mutual conversion (**Figure 1b and 1c**).

In this paper, we showed that random mutations not only rapidly establish positive or negative mutually regulatory relationships between two unrelated fused proteins regardless of the linker length or linkage format, but also frequently reverse their regulatory logic. This surprising ability of fusion proteins to evolve reciprocal regulation is likely to be closely linked to the emergence of BIF through the destabilizing effect of random mutations. Considering this remarkable frequency of appearance, binding-induced stabilization could become one of the major operating principles of natural protein switches. Indeed, some transcription factors, such as TraR regulating the quorum sensing of *Agrobacterium tumefaciens*^44^, are known to be unable to fold by themselves. Recent studies on the tetracycline (Tc) -induced transcription repressor TetR, which had been described as an allosteric switch, are revealing that it is at least partially operated through ligand-induced folding and/or MEF. One model has been proposed in which Tc binding reinforces TetR with an “DNA-unbound” state, thereby reducing the capacity of induced fitting of TetR to its operator DNA^45^.

Additionally, another model has been proposed in which the stabilization of the LBD by Tc binding leads to the destabilization of the DBD, thereby leading to the detachment of TetR from its operator^46^. Moreover, the reversal of behavior has been reported for several transcription factors, and many of these reversals are suspected to be driven by ligand-induced folding^45,47,48^. Taken together, our results have demonstrated that the destabilizing effect of random mutations, which has long been regarded as the greatest obstacle to evolving protein function^49^, is an extremely useful evolutionary resource for synthetic biologists, at least in the context of designing, improving, and modifying the specifications and/or their operational logics of molecular switches.

## Materials and Methods

### Bacterial strains, media, and growth conditions

*Escherichia coli* strain XL10-Gold (Agilent Technology, Inc., Santa Clara, CA) was used for standard molecular cloning. *E. coli* strain JW0063 without a kanamycin resistance cassette was used for library construction^50^, directed evolution, and all functional assays. The genotypes are provided in **Table S6**. For all experiments, *E. coli* strains were grown at 37°C with the appropriate antibiotics at the following concentrations in LB liquid medium (2% w/v Lennox LB; Nacalai Tesque, Kyoto, Japan) or LB agar plates (2% w/v Lennox LB; Nacalai Tesque, 1.5% (w/v) agar; Nacalai Tesque), with 100 µg/mL ampicillin (Sigma-Aldrich, Inc, St. Lous, MO), 30 µg/mL chloramphenicol (Nacalai Tesque), and 30 µg/mL streptomycin (Nacalai Tesque), unless otherwise stated. The stock solutions (1 M) of L-arabinose (Tokyo Chemical Industry Co., Ltd., Tokyo, Japan) and D-fucose (Tokyo Chemical Industry Co., Ltd.) were prepared by dissolving appropriate amounts of the compounds in deionized water and filter-sterilizing them through a 0.2 µm cellulose acetate filter. The plasmid and primer lists are provided in **Tables S7** and **S8**, respectively.

### AraC::CAT library construction with randomized linkers

The *cat* gene was amplified using partially randomized primers (**Table S8**) and KOD Plus DNA Polymerase (TOYOBO, Osaka, Japan). The plasmid backbone containing the region of the split *araC* genes was amplified by using the appropriate primers and KOD Plus. These amplicons were purified using a FastGene Gel/PCR Extraction Kit (Nippon Genetics, Tokyo, Japan) and then subjected to Golden Gate assembly. After these products were purified and introduced into JW0063 by electroporation, the library plasmids were obtained from transformants by miniprep (FastGene PlasmidMini; Nippon Genetics). The size of each library was estimated to be approximately 10^5^ colonies.

### Whole-gene mutagenesis

Initially, high-fidelity PCR was performed using KOD Plus to amplify the *araC::cat* region or the mutants on the pHRA-based vector. The amplicons were purified and subjected to error-prone PCR under the following conditions: 5 U Taq DNA polymerase (New England Biolabs, MA, USA), 200 μM of each deoxynucleoside triphosphate, 2 mM MgCl_2_, and 50 μM MnCl_2_. The amplification factor was about 10^3^. The purified error-prone PCR products were digested using BamHI-HF (New England Biolabs, MA, USA) and HindIII-HF (New England Biolabs) and then ligated into a pHRA-based vector using T4 DNA ligase (New England Biolabs). The ligation mixture was purified and then transformed into JW0063 by electroporation. DNA from the transformants was extracted via miniprep to obtain the library plasmid (library size is approximately 10^5^).

### Elimination of dead mutants and library evaluation

To eliminate the *araC::cat* genes that encoded dead mutants and cloning artifacts, JW0063 harboring pAraC::CAT libraries were inoculated on solid LB medium containing ampicillin and chloramphenicol at 37 °C for 12 h. To evaluate CAT activity, 88 randomly selected colonies were grown overnight in LB medium. Then, 4 µL of the preculture was inoculated into 400 µL of LB medium containing ampicillin, various concentrations of chloramphenicol, and 0 or 10 mM of D-fucose in a 96-deep well plate. After shaking at 37°C for 12 h, cell densities were measured at 595 nm (OD_595_) using FilterMax F5 (Molecular Devices). The FD score is defined below:

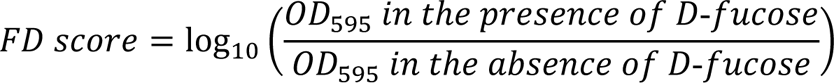

where the concentration of D-fucose is 10 mM. Matplotlib, a module for visualizing graphs in Python, was used for data analysis.

### Gene expression analysis using fluorescent proteins as AraC function

To quantify the AraC function, JW0063 harboring plasmids encoding the P_BAD7_-sfgfp-HSVtk-aph and *araC::cat* genes were incubated on solid LB medium. The overnight culture from a single colony was inoculated with 1% culture medium in 400 µL of liquid LB medium containing appropriate antibiotics and with or without 10 mM of D-fucose. in 96-deep well plates. After incubation for 12 h at 37°C, these cultures were diluted 10-fold in 20 µL culture medium-saline (0.9% (w/v) NaCl; Nacalai Tesque) and placed in 96-well shallow plates. Cell density (OD_595_) and green fluorescence (ex 485 nm, ex 535 nm) were measured using a FilterMax F5. Fluorescence values were normalized to OD_595_.

Single-cell analysis by flow cytometry was performed using a MACS Quant VYB (Miltenyi Biotech, Bergisch-Gladbach, Germany). The cell cultures described above were diluted at approximately 1:100, and 50,000 cells were analyzed. The measurements were performed at an FSC voltage of 320 V, an SSC voltage of 230 V, and a B1 laser (excited at 488 nm and emitted at 525/50 nm) voltage of 340 V. Data were analyzed using a MACSQuant analyzer (Miltenyi Biotech, Bergisch-Gladbach, Germany).

### Generating model structures

Model structures of the fusion proteins were generated as follows. The accessible surface area of AraC was calculated by PyMOL to determine the potential positions for CAT insertion. Using MODELLER, the CAT monomer was inserted into the AraC loop via linker peptides (N: GGSGGSGGS, C; ACGSGSGSG). For the calculation, the PDB entries of 2arc, 2k9s, and 3u9f were adopted as AraC’s LBD, AraC’s DBD, and CAT, respectively.

### FoldX calculation

To remove unfavorable torsion angles from the AraC::CAT model structure predicted by using AlphaFold2, van der Waals collisions, and total energies from the predicted model, the side chains of the model structure were rearranged using the RepairPDB command in FoldX to generate stabilized structures. The free energy change (ΔΔG) between wild-type and mutant was predicted using the BuildModel command with the following settings: ionStrength = 0.05, pH = 7, temperature = 298 K, vdwDesign = 2, moveNeighbours = true, number of runs = 3. The mean value was used for analysis.

## Author contributions

All authors designed the research. YY and MT performed all experiments with assistance from YK and DU. All authors wrote the manuscript.

## Acknowledgements

This work was conducted as a part of the Waseda Research Institute for Science and Engineering project “Experimental Evolution of Genetic Devices (22P04), and project, JPNP20011, commissioned by the New Energy and Industrial Technology Development Organization (NEDO).” DU was supported by JSPS KAKENHI Grant Numbers 21H01721, 18H01791, and 16H06450, The Salt Science Research Foundation (No. 2201), Futaba Electronics Memorial Foundation, and Kioxia Corporation.

