## Supporting Information for "Fabrication, evolution, and mutual conversion of D-fucose-activatable and -repressible acetyltransferase upon mutations"

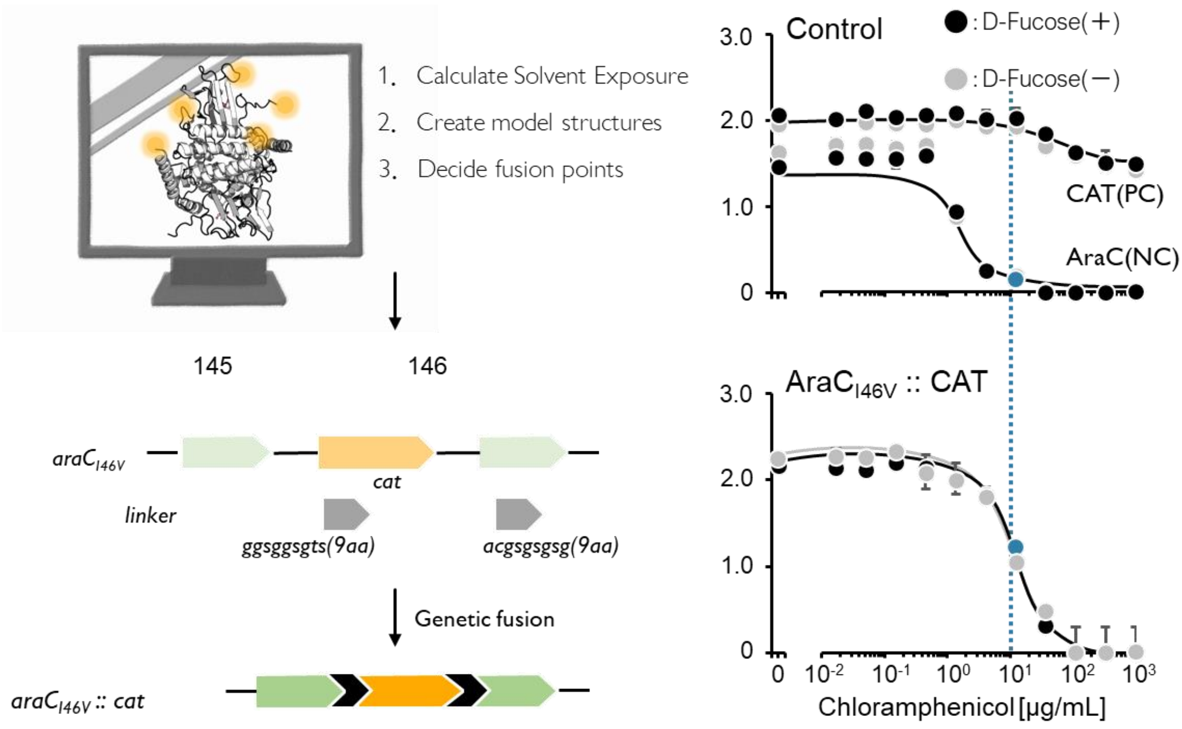


**Figure S1. CAT functions of AraC$_{I46V}$-CAT (GS linker).**

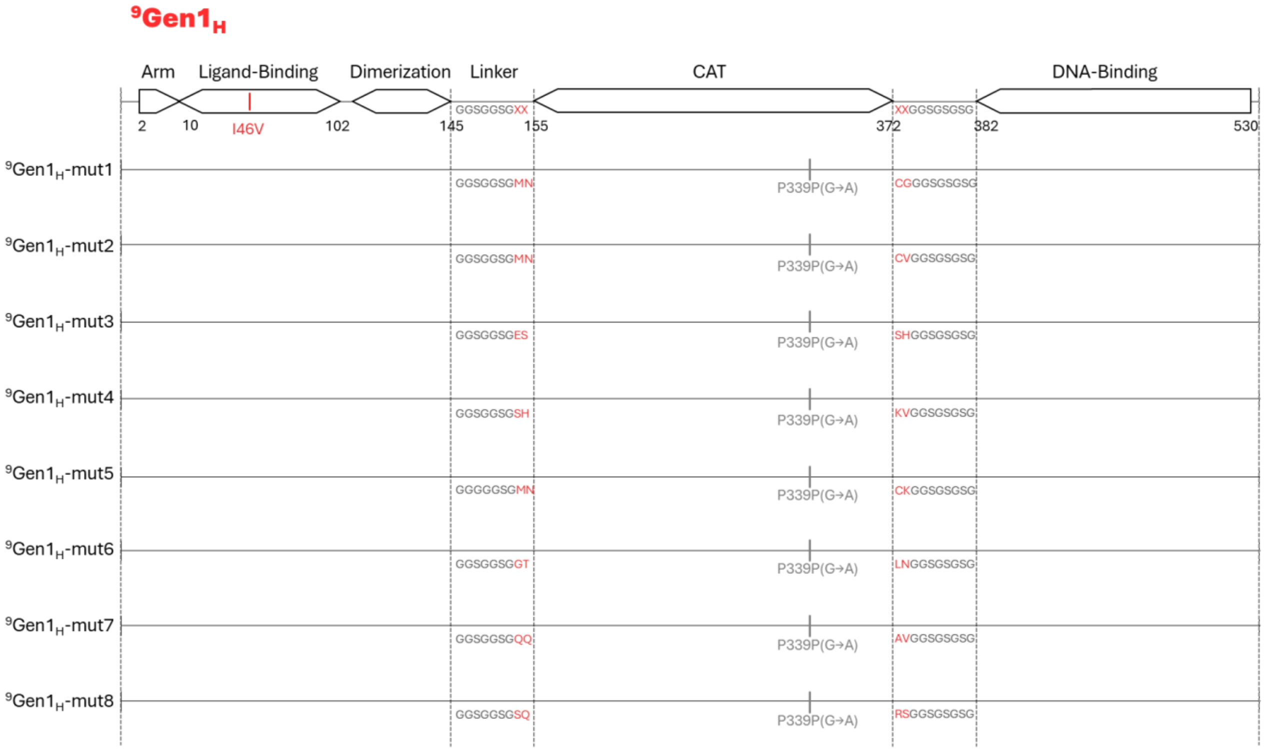


**Figure S2. Sequences of the high-FD variants $^{9}$Gen1$_{H}$.**

Arranged from top to bottom in order of FD score. Red lines are introduced mutations, grey lines are synonymous mutations (base changes in brackets). Red letters in the linkers indicate the two randomized amino acids.

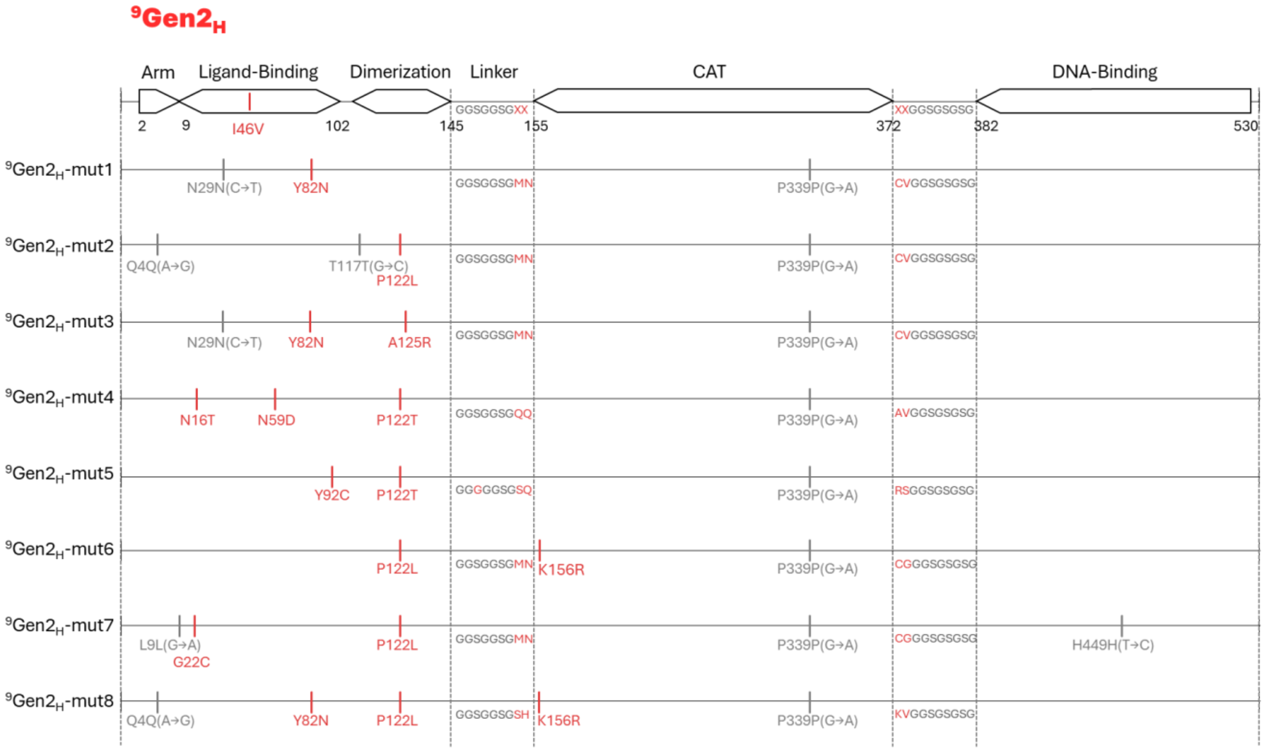


**Figure S3. Sequences of the high-FD variants ⁹Gen2$_H$.**

Arranged from top to bottom in order of FD score. Red lines are introduced mutations, grey lines are synonymous mutations (base changes in brackets). Red letters in the linkers indicate the two randomized amino acids.

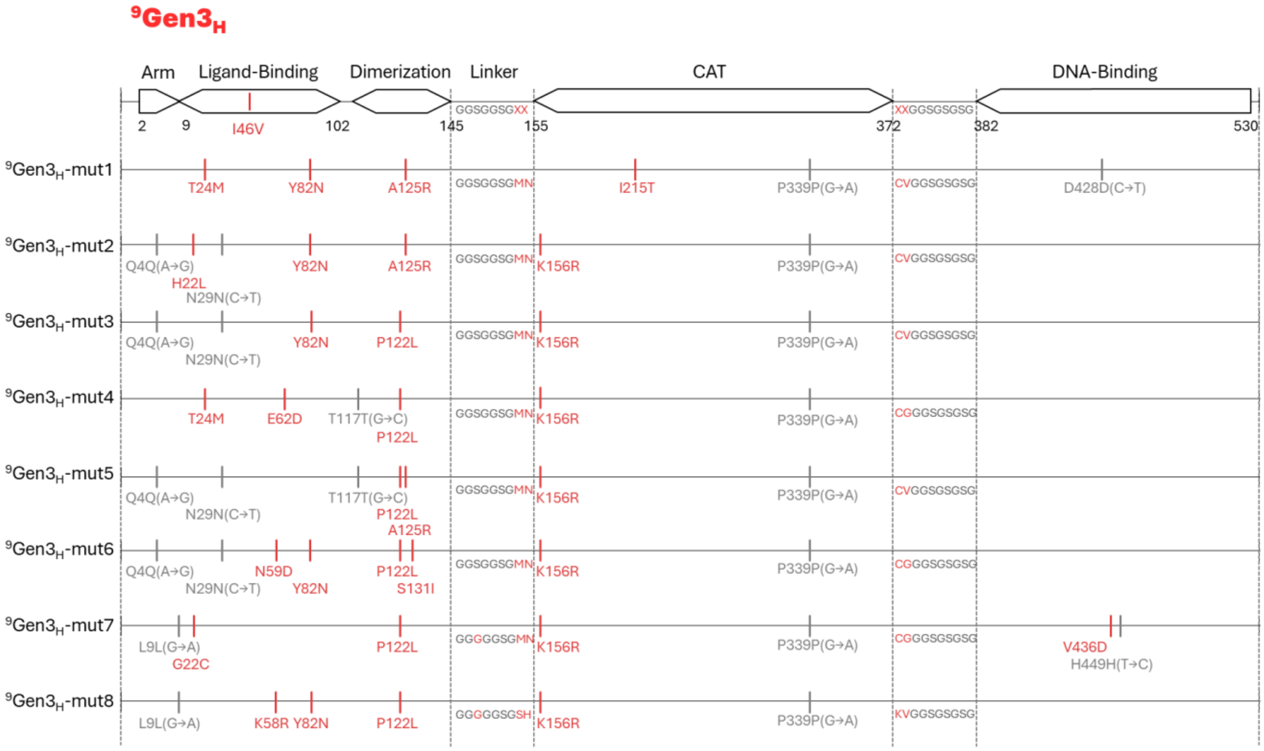


**Figure S4. Sequences of the high-FD variants $^{9}Gen3_{H}$.**

Arranged from top to bottom in order of FD score. Red lines are introduced mutations, grey lines are synonymous mutations (base changes in brackets). Red letters in the linkers indicate the two randomized amino acids.

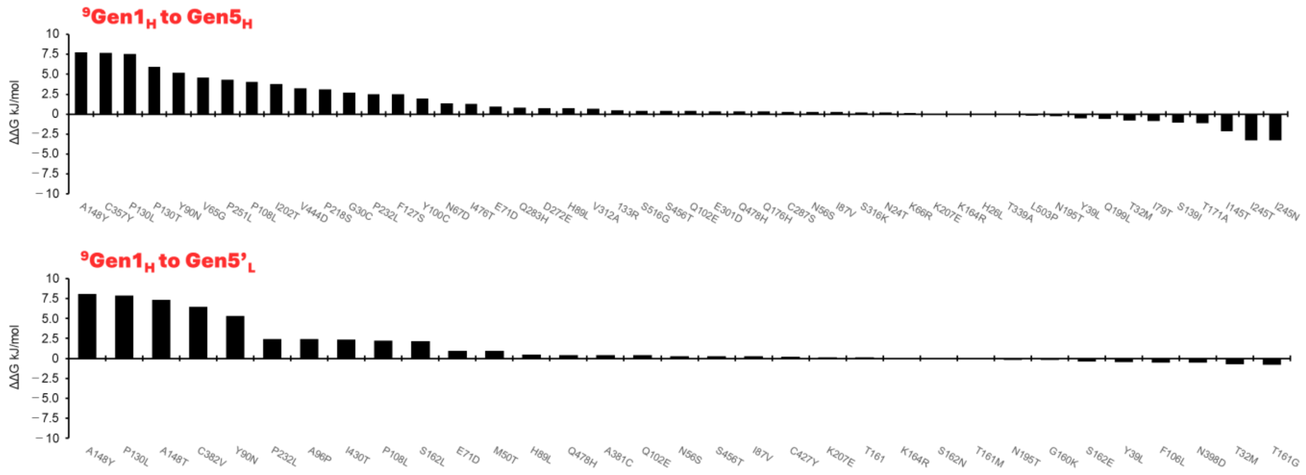


**Figure S5. Calculated ΔΔG for each of the mutations found in the AraC::CAT variants with (9 + 9)-linkers ($^{9}$Gen1, $^{9}$Gen2, $^{9}$Gen3, $^{9}$Gen4 $^{9}$Gen5, or $^{5}$Gen5').**

The *ΔΔ*G of each of mutations found in mutants with a high FD scores $^{9}$Gen1 to Gen5 were calculated using Fold-X. The vertical axis represents *ΔΔ*G (kJ/mol) calculated by FoldX.

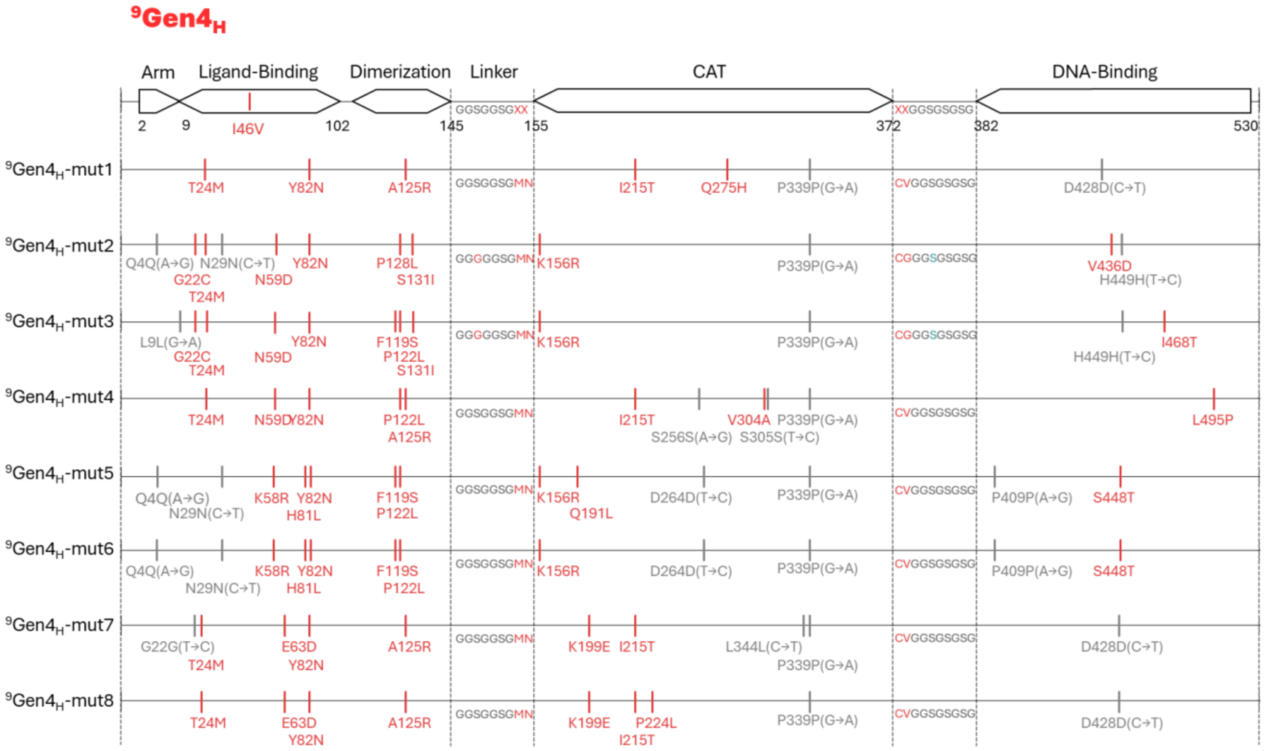


**Figure S6. Sequences of the high-FD variants ⁹Gen4H.**

Arranged from top to bottom in order of FD score. Red lines are introduced mutations, grey lines are synonymous mutations (base changes in brackets). Red letters in the linkers indicate the two randomized amino acids.

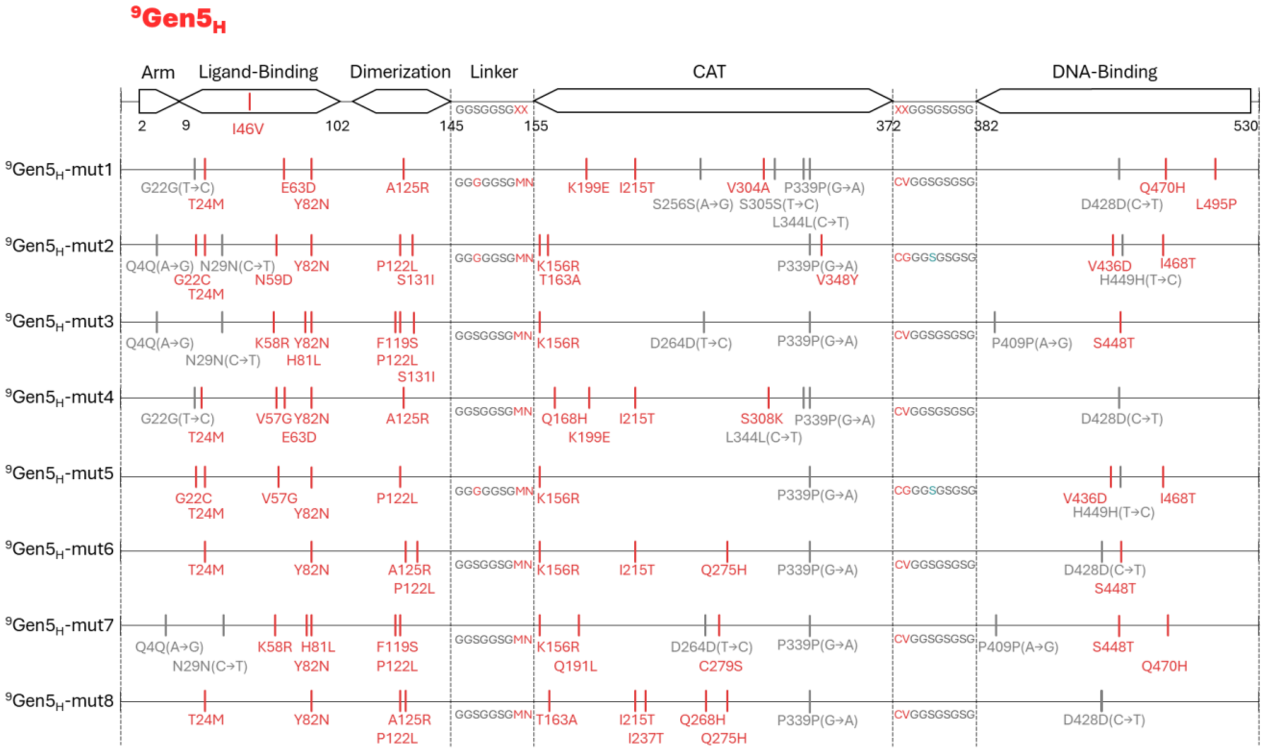


**Figure S7. Sequences of the high-FD variants $^{9}$Gen5$_{H}$.**

Arranged from top to bottom in order of FD score. Red lines are introduced mutations, grey lines are synonymous mutations (base changes in brackets). Red letters in the linkers indicate the two randomized amino acids.

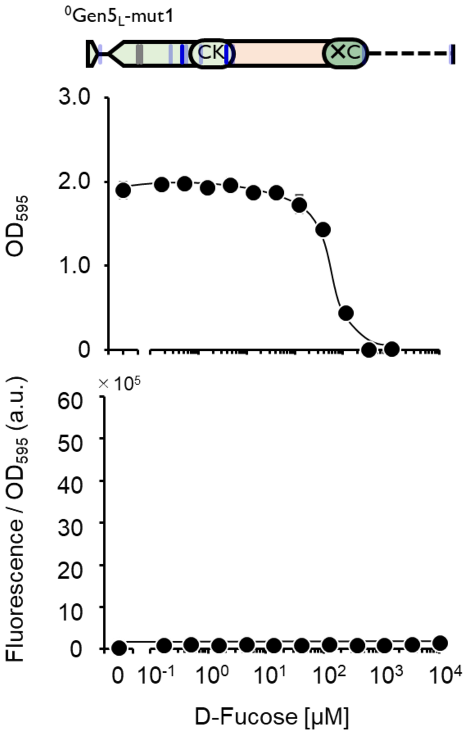


**Figure S8. D-Fucose-dependent curves of CAT and AraC function of $^{0}$Gen5$_L$-1 (with one terminal codon).**

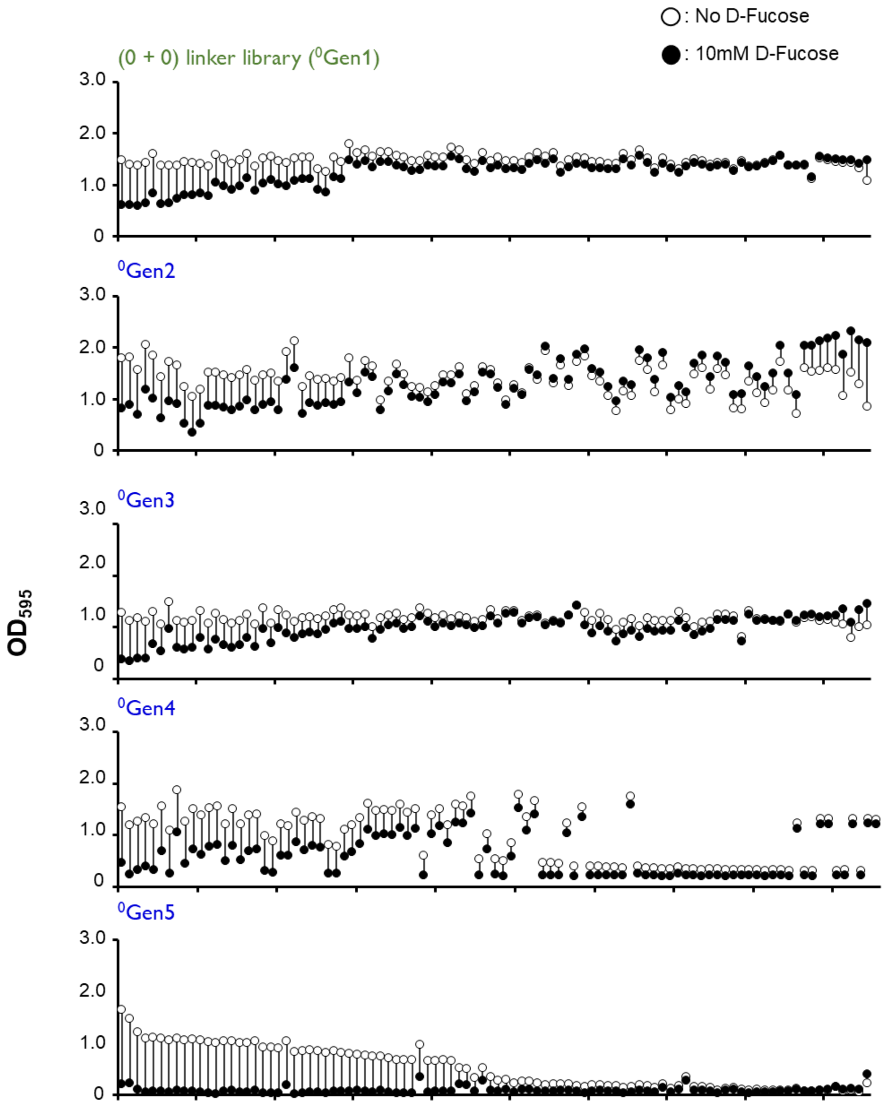


**Figure S9. Functional distribution of (0 + 0) linker variants.**

Clones selected in the first screening were randomly selected and plotted for $OD_{595}$ after overnight incubation in liquid medium with (black circles) or without (unfilled) D-fucose. The length of the line represents the difference in OD595 between the two conditions.

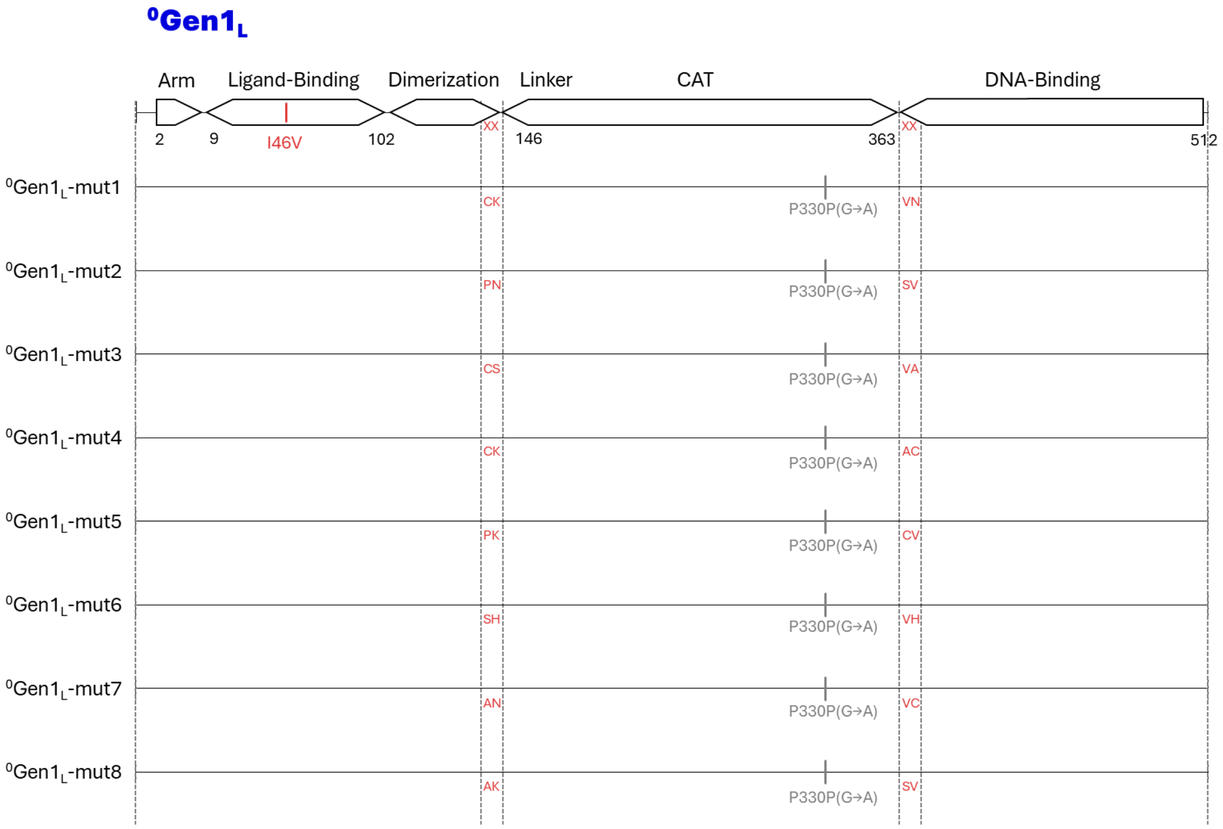


**Figure S10. Sequences of the low-FD variants $^{0}$Gen1$_{L}$.**

Arranged from top to bottom in order of FD score. Grey lines are synonymous mutations (base changes shown in brackets). The red letters in the linkers indicate the two randomized amino acids.

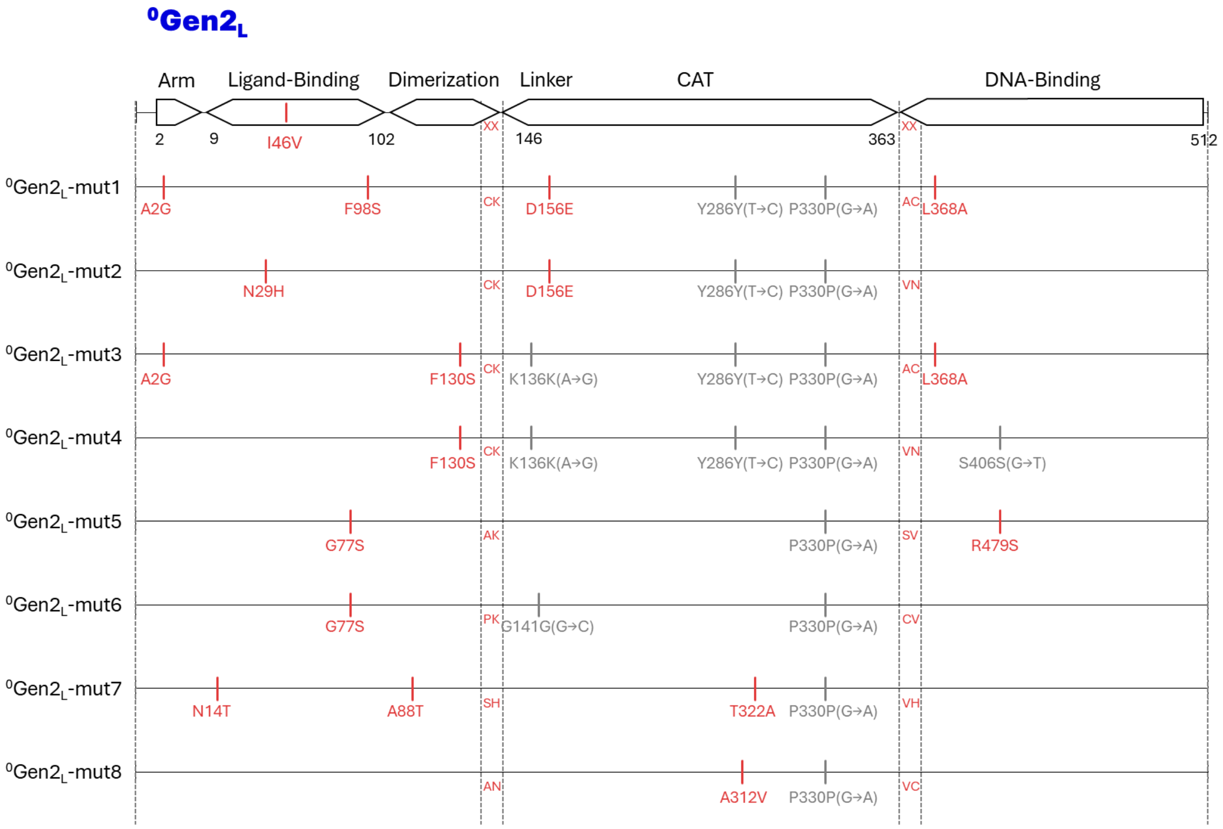


**Figure S11. Sequences of the low-FD variants $^{0}$Gen1$_{L}$.**

Arranged from top to bottom in order of FD score. Red lines are introduced mutations, and grey lines are synonymous mutations (base changes in brackets). Red letters in the linkers indicate the two randomized amino acids.

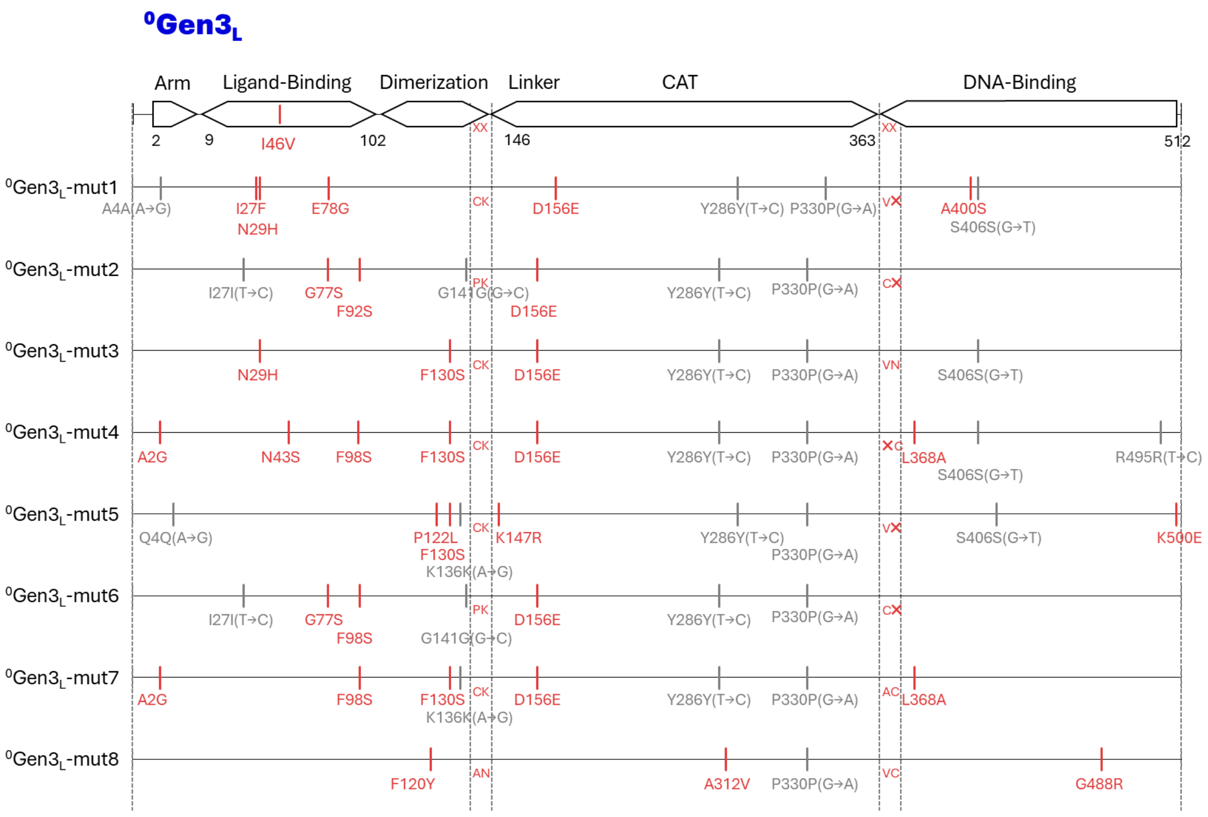


**Figure S12. Sequences of the low-FD variants $^{0}$Gen3$_{L}$.**

Arranged from top to bottom in order of FD score. Red lines are introduced mutations, and grey lines are synonymous mutations (base changes in brackets). Red letters in the linkers indicate the two randomized amino acids.

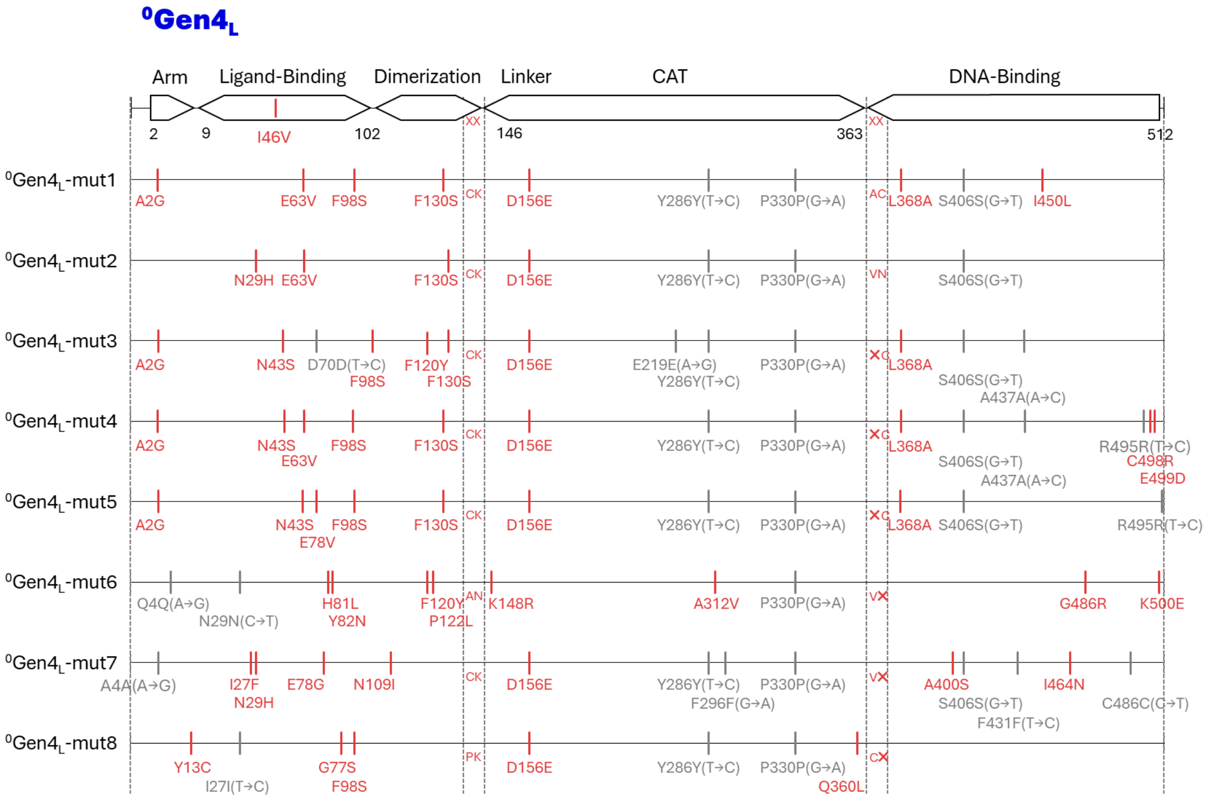


**Figure S13. Sequences of the low-FD variants $^{0}$Gen4$_{L}$.**

Arranged from top to bottom in order of FD score. Red lines are introduced mutations, and grey lines are synonymous mutations (base changes in brackets). Red letters in the linkers indicate the two randomized amino acids.

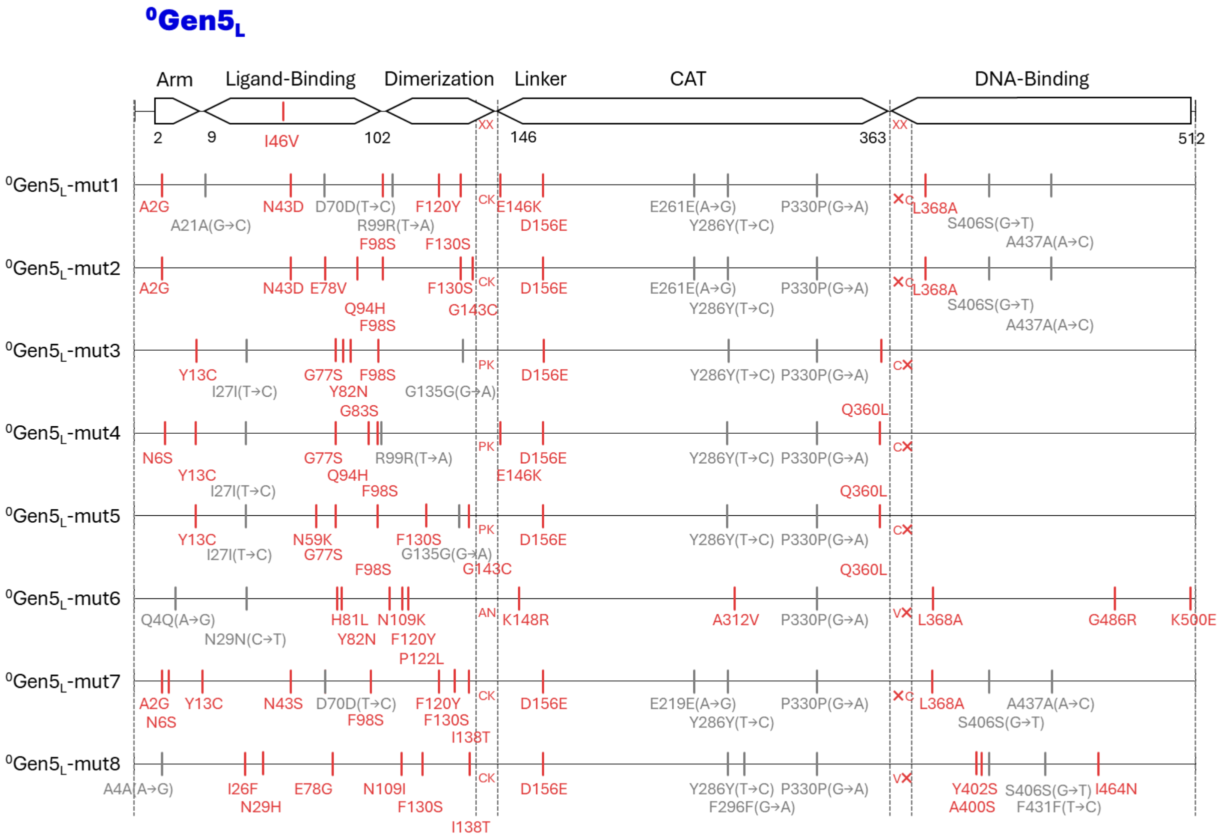


**Figure S14. Sequences of the low-FD variants $^{0}$Gen5$_{L}$.**

Arranged from top to bottom in order of FD score. Red lines are introduced mutations, and grey lines are synonymous mutations (base changes shown in brackets). Red letters in the linkers indicate the two randomized amino acids.

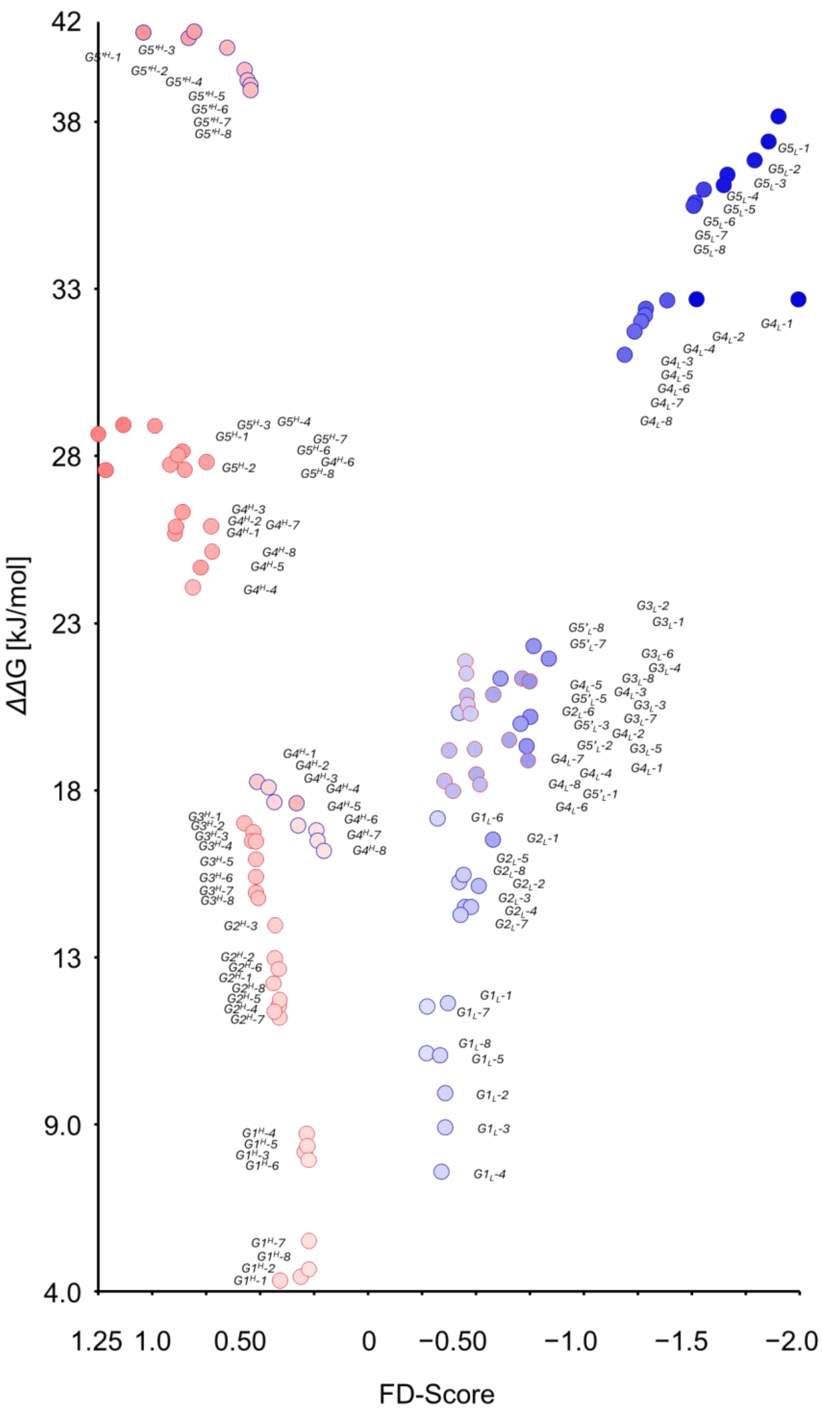


**Figure S15. Relationship between calculated ΔΔG values and FD scores for all the mutants analyzed in two lineages.**

The *ΔΔ*G, which is the sum of all mutations introduced throughout the gene for each mutant, was calculated using FoldX using structural model created by Alphafold. The frame lines of the plots are painted red for the variants with (9+9) linkers, and are blue for those with (0+0) linkers.

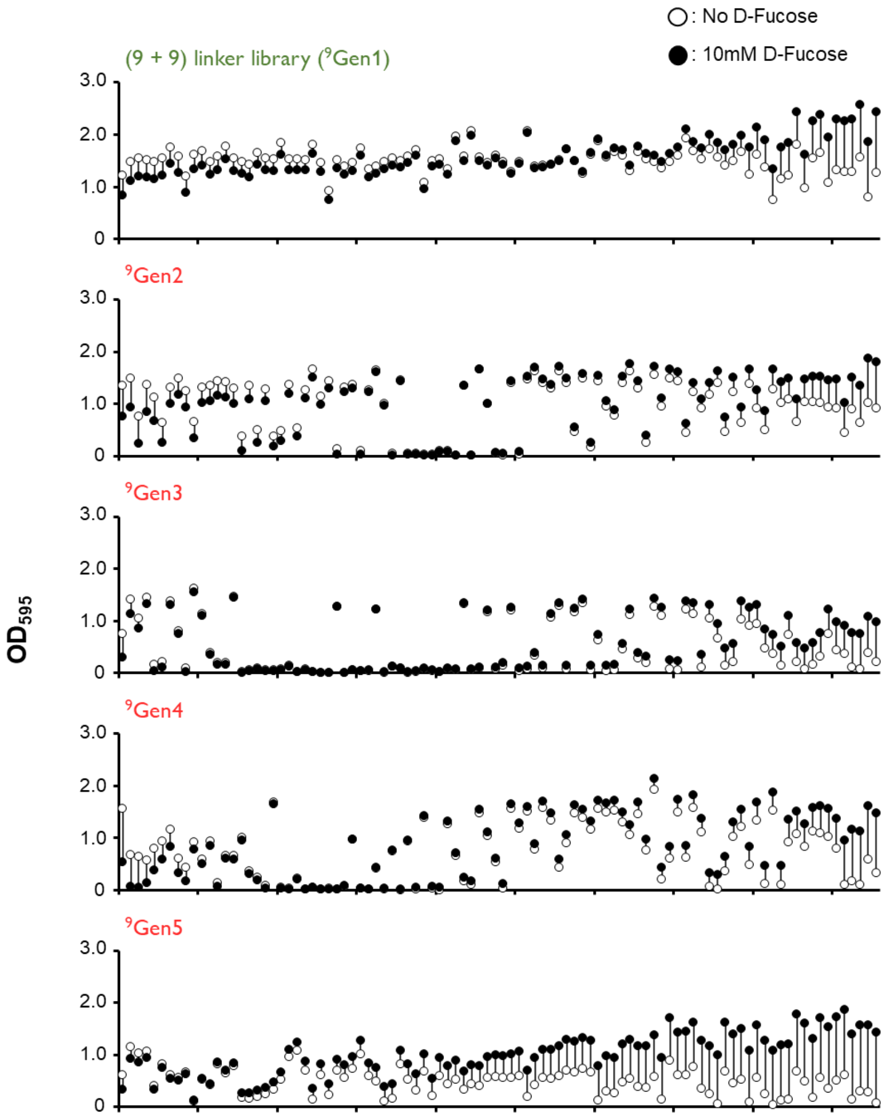


**Figure S16. Functional distribution of (9 + 9) linker variants.**

Clones selected in the first screening were randomly selected and plotted for $OD_{595}$ after overnight incubation in liquid medium with (black circles) or without (unfilled) D-fucose. The length of the line represents the difference in OD595 between the two conditions.

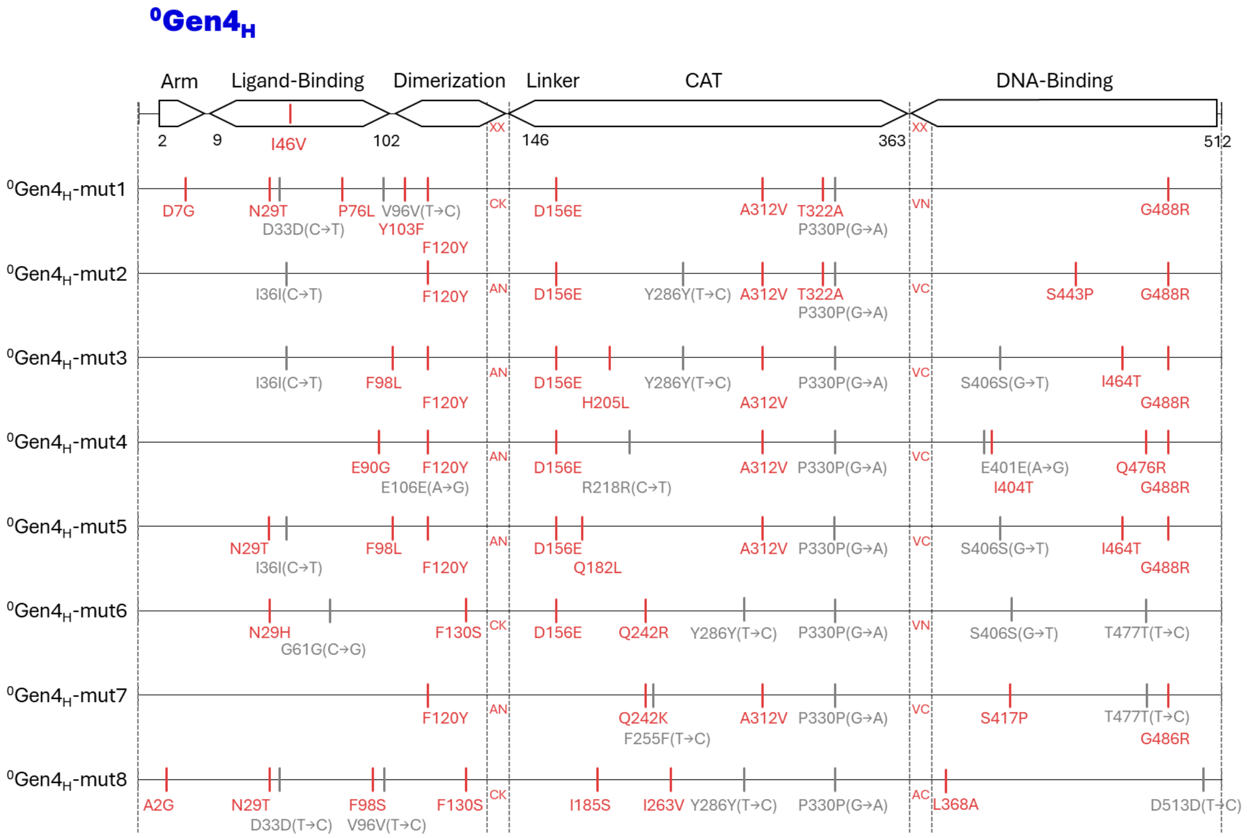


**Figure S17. Sequences of the high-FD variants $^{0}$Gen4$_H$.**

Arranged from top to bottom in order of FD score. Red lines are introduced mutations, and grey lines are synonymous mutations (base changes in brackets). Red letters in the linkers indicate the two randomized amino acids.

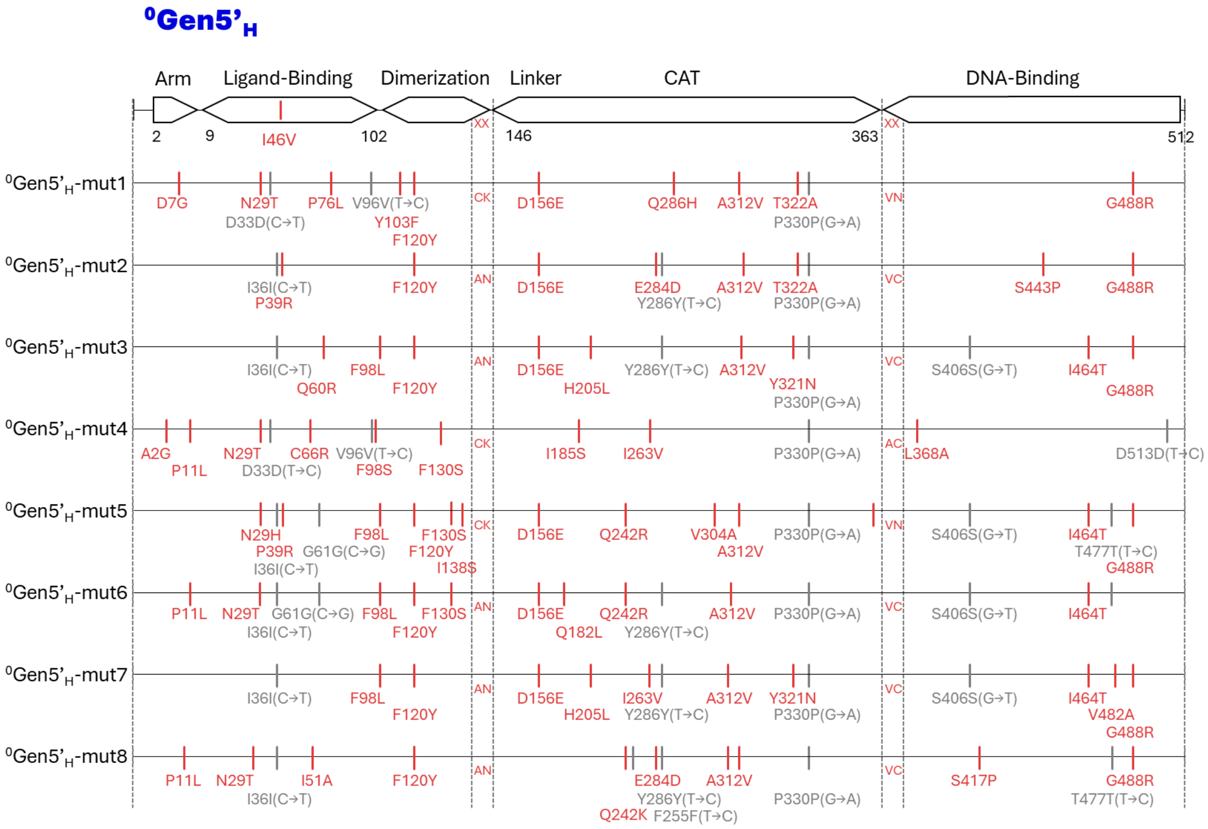


**Figure S18. Sequences of the reversed, high-FD variants $^{0}$Gen5'$_{H}$.**

Arranged from top to bottom in order of FD score. Red lines are introduced mutations, and grey lines are synonymous mutations (base changes in brackets). Red letters in the linkers indicate the two randomized amino acids.

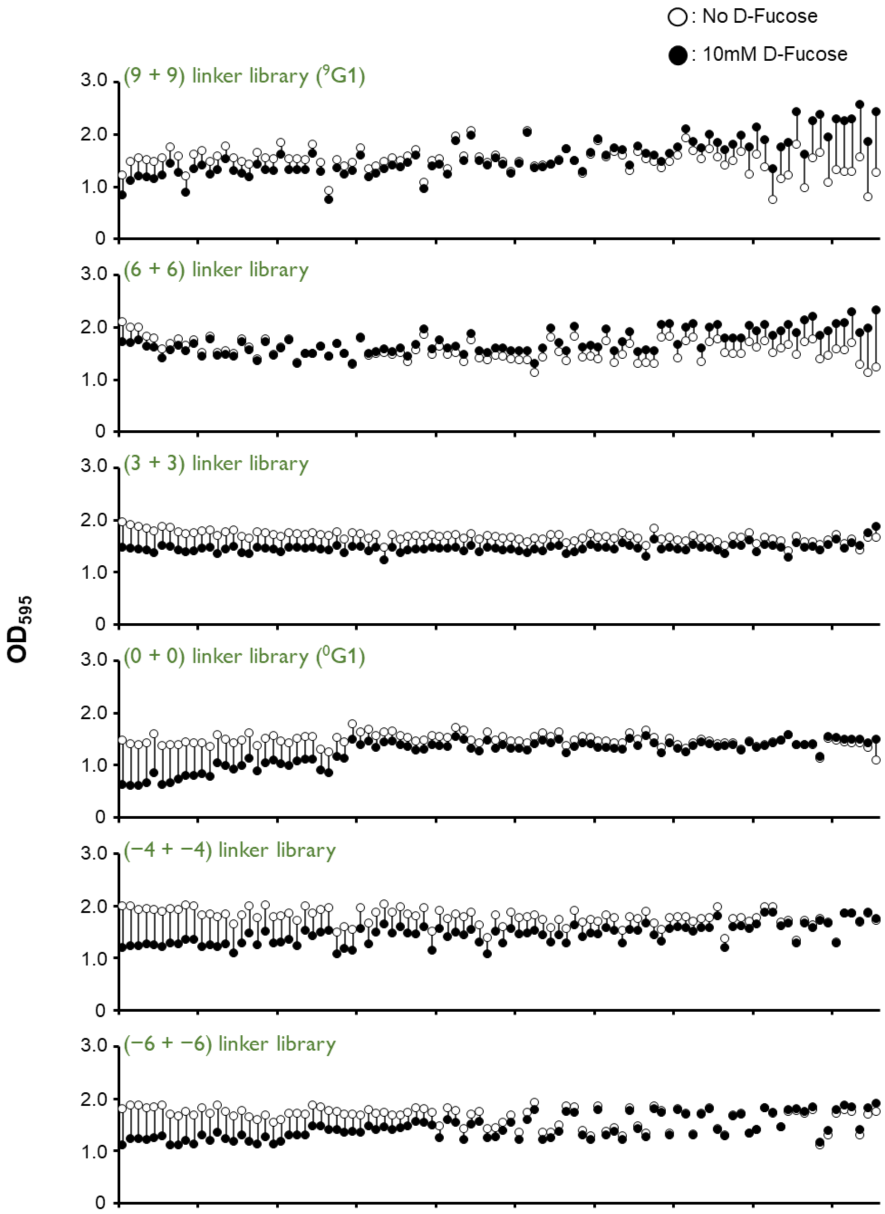


**Figure S19. Functional distribution of Gen1 variants with different linker sizes.**

Clones selected in the first screening were randomly selected and plotted for $OD_{595}$ after overnight incubation in liquid medium with (black circles) or without (unfilled) D-fucose. The length of the line represents the difference in OD595 between the two conditions.

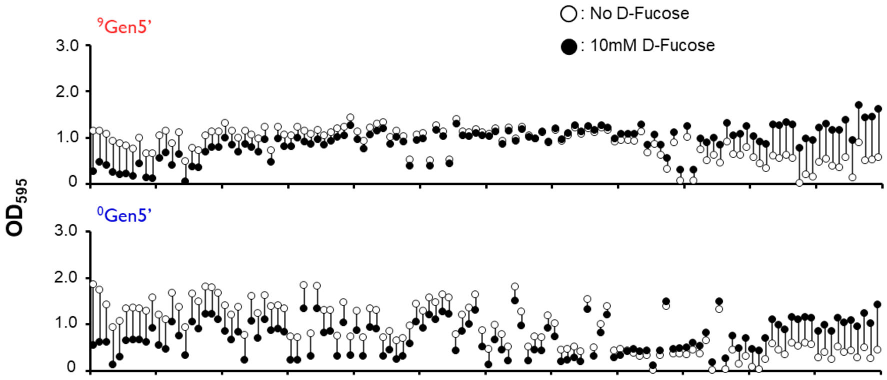


**Figure S20. Functional distribution of Gen5' variants with reversed phenotypes.**

Clones selected in the first screening were randomly selected and plotted for $OD_{595}$ after overnight incubation in liquid medium with (black circles) or without (unfilled) D-fucose. The length of the line represents the difference in OD595 between the two conditions.

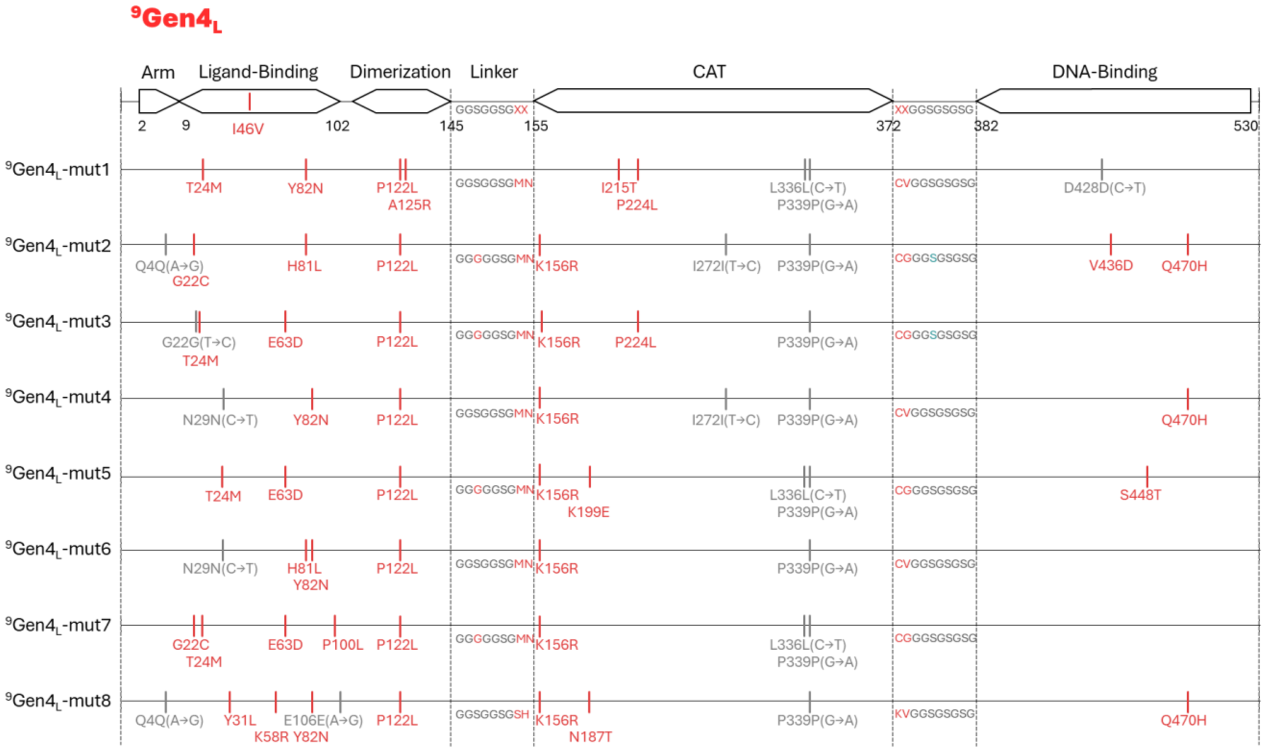


**Figure S21. Sequences of the low-FD variants $^{9}Gen4_{L}$.**

Arranged from top to bottom in order of FD score. Red lines are introduced mutations, and grey lines are synonymous mutations (base changes in brackets). Red letters in the linkers indicate the two randomized amino acids.

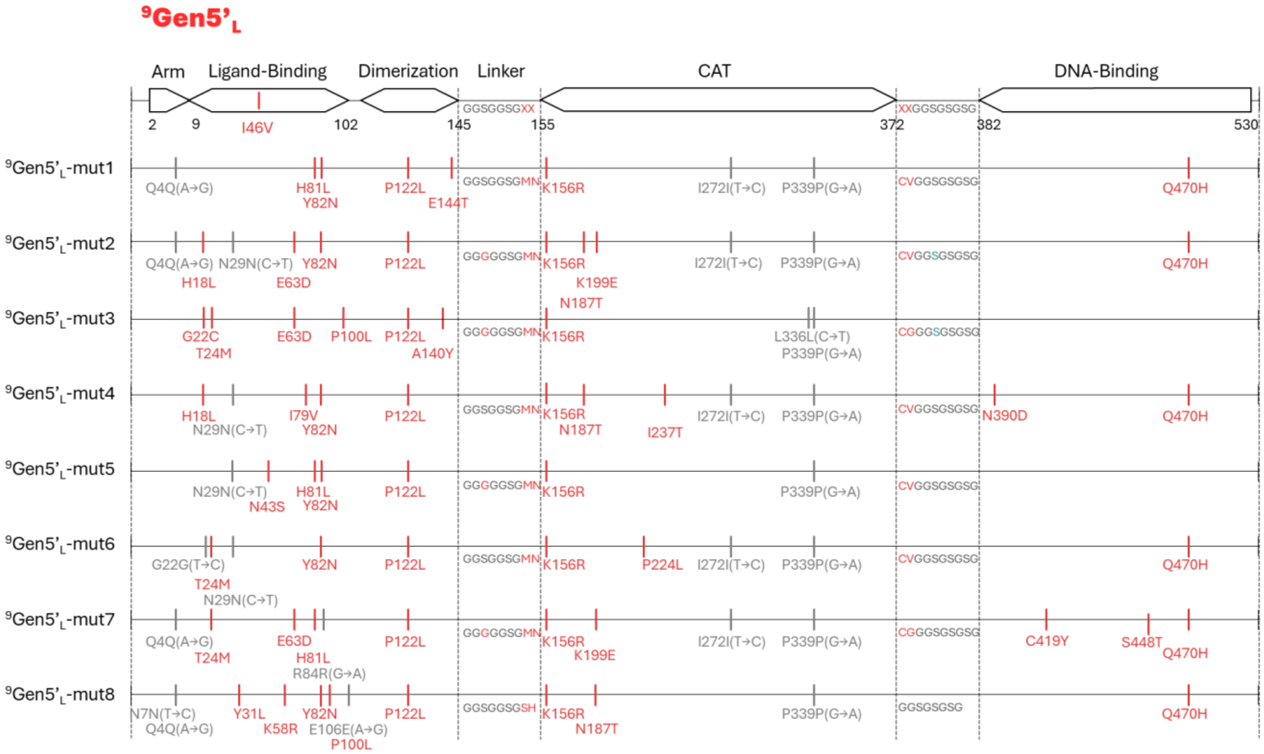


**Figure S22. Sequences of the low-FD variants [9]Gen5'$_L$.**

Arranged from top to bottom in order of FD score. Red lines are introduced mutations, and grey lines are synonymous mutations (base changes in brackets). Red letters in the linkers indicate the two randomized amino acids.

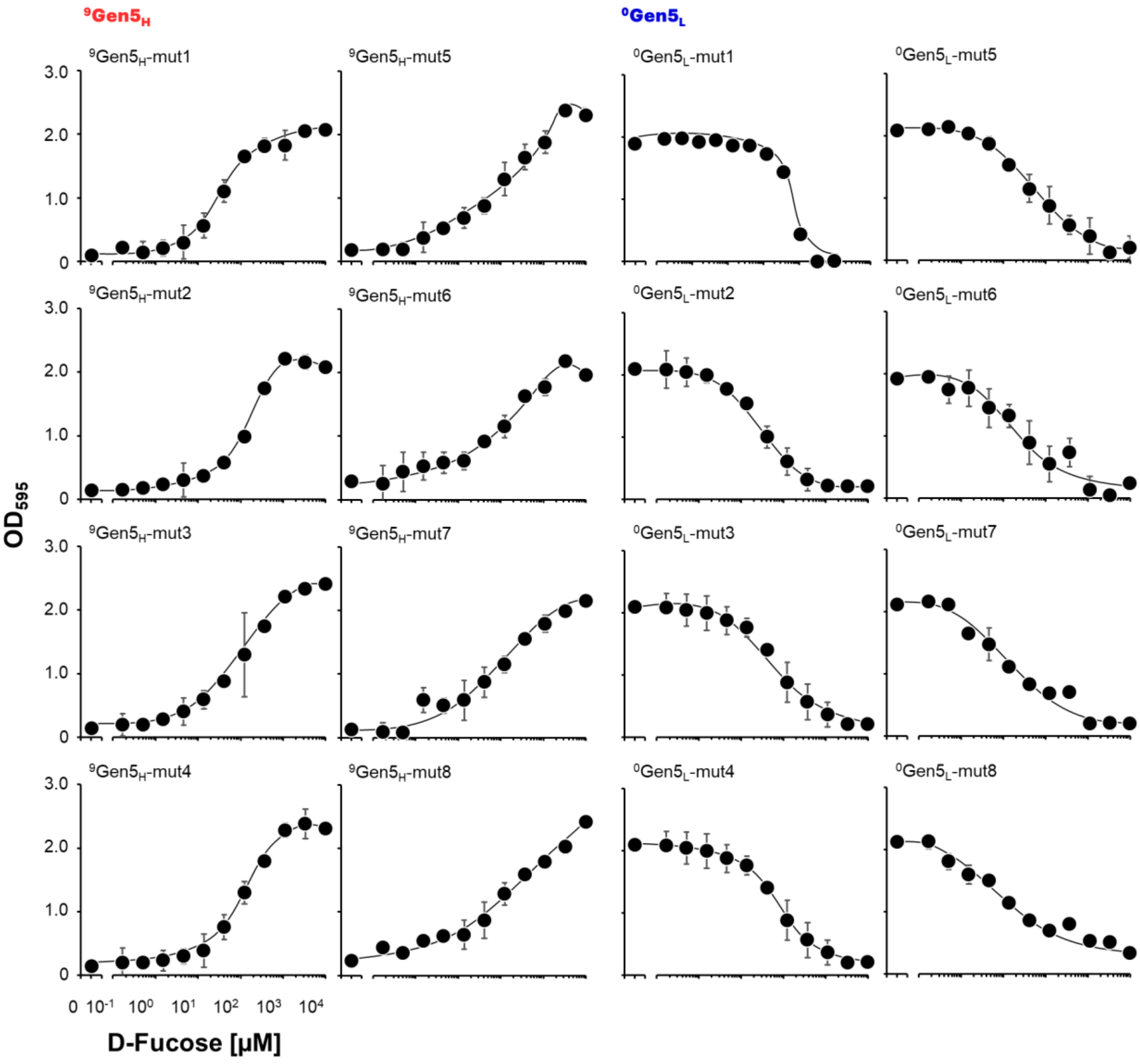


**Figure S23. Dose dependency of CAT activity of the best switchers obtained at generation-5 in the two lineages on D-Fucose concentration.**

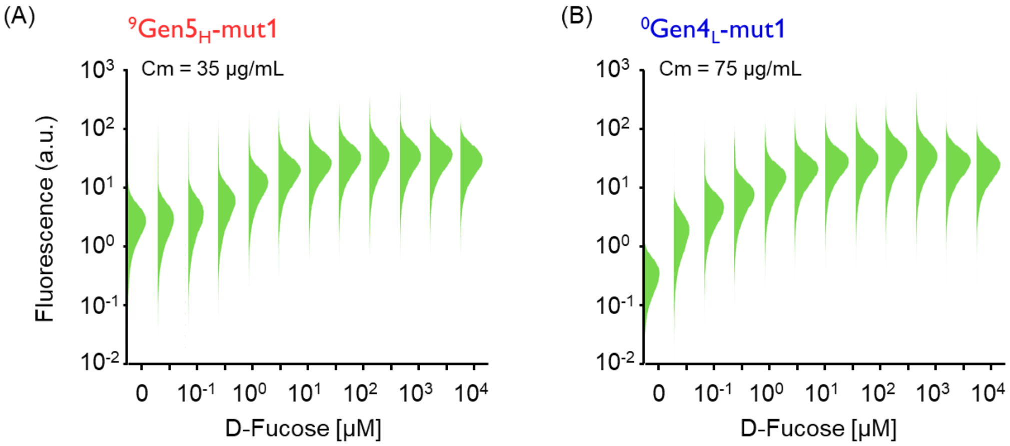


**Figure S24. D-Fucose dose dependent changes in CAT function under Cm condition.**

Fluorescence intensity was measured by single-cell analysis with flow cytometry. The average of the fluorescence values measured for 50,000 *E. coli* was used as the data for each concentration.

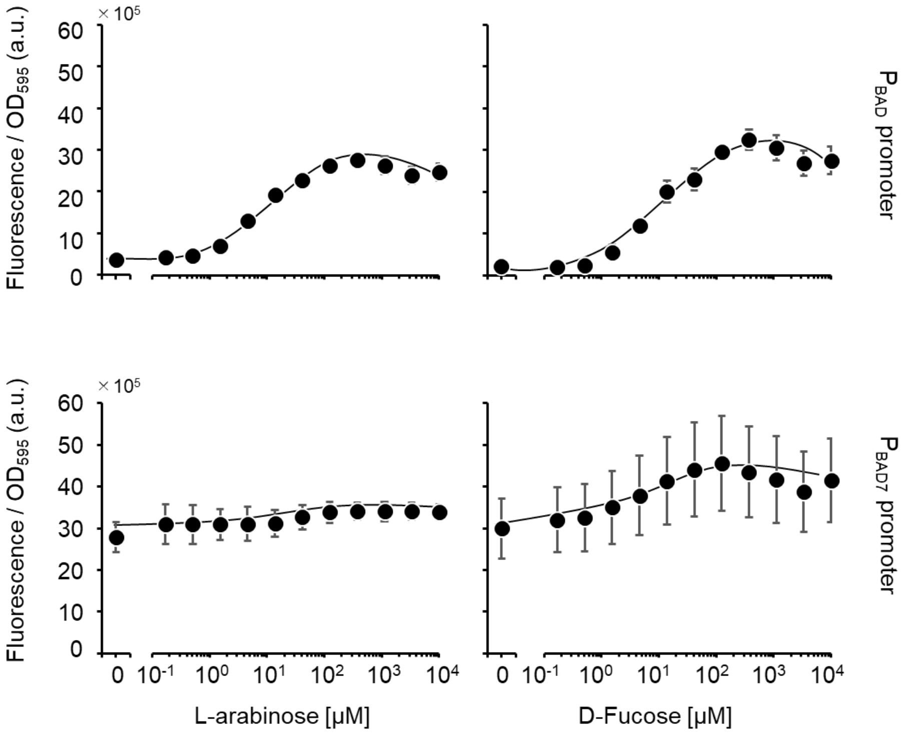


**Figure S25. L-arabinose and D-Fucose dependent behaviors of prototype AraC$_{I46V}$-CAT (with GS linker) on P$_{BAD}$ and P$_{BAD7}$.**

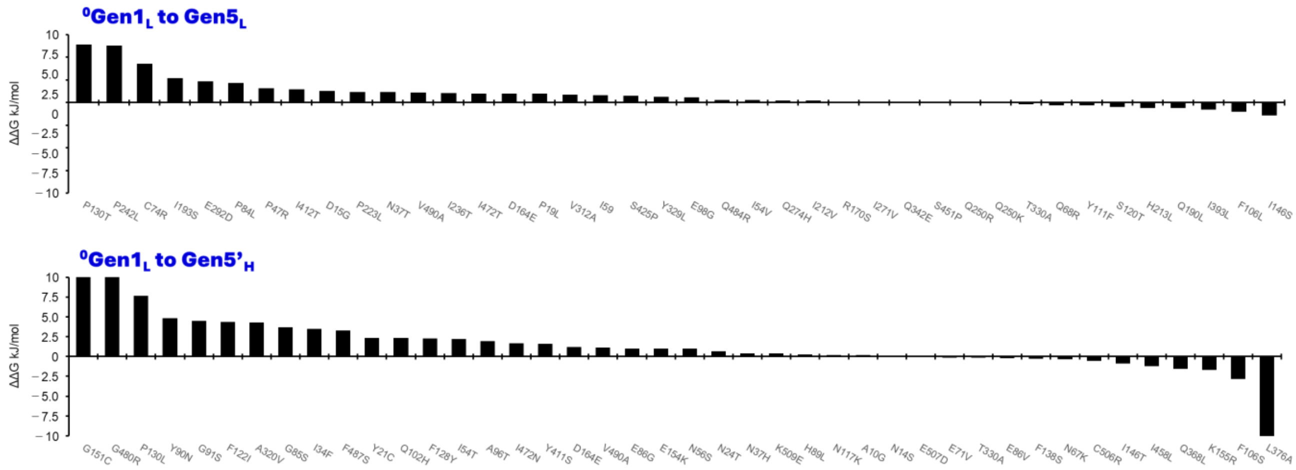


**Figure S26. Calculation of ΔΔG for each of the mutations introduced in the (0 + 0) (Gen1 to 5 or 5') by FoldX.**

The format of the figure is the same as in **Figure S5.**

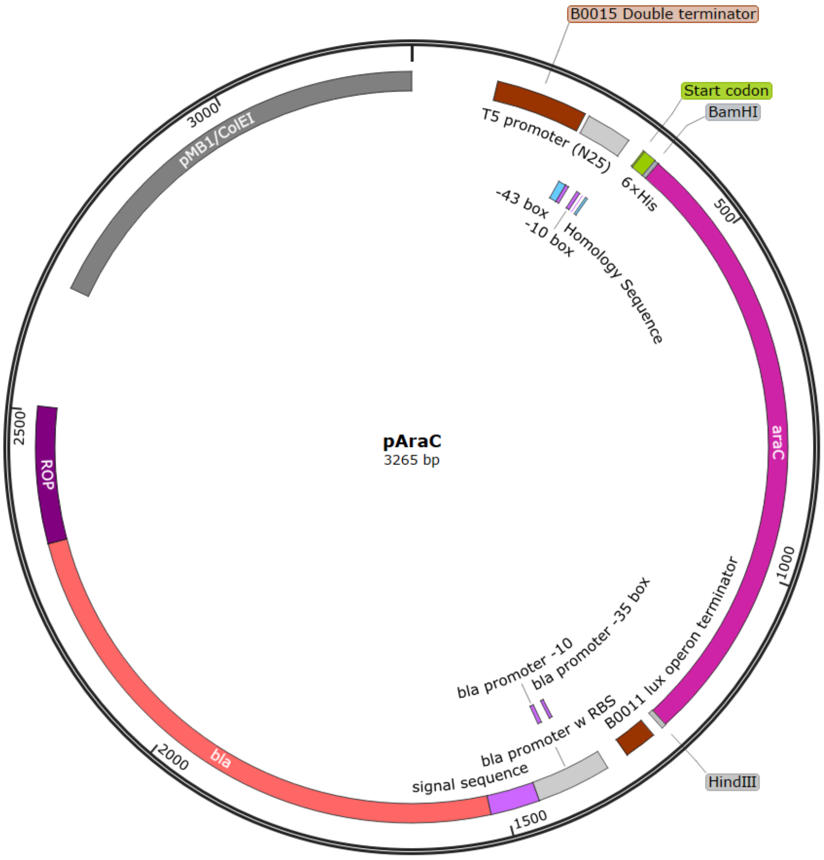


AACGCCAGCAACGCGGCCTTTTTACGGTTCCTGGCCTTTTGCTGGCCTTTTGCTCACATGTTCTTTCCTGCGTTATC
CCCTGATTCTGTGGATAACCGTATTACCGCCTTTGAGTGAGCTCCAGGCATCAAATAAAACGAAAGGCTCAGTCGAA
AGACTGGGCCTTTCGTTTTATCTGTTGTTTGTCGGTGAACGCTCTCTACTAGAGTCACACTGGCTCACCTTCGGGTG
GGCCTTTCTGCGTTTATAGGTACCAAGAATCATAAAAAATTTATTTGCTTTCAGGAAAATTTTTCTGTATAATAGAT
TCATAAATTTGAGGGCCCCCCTCCTTCAGGTCTAAAAGTCGCAATGTTGGTTTTGACGATCAACTCTATTTCTCGCG
AGTATTTAAAAAATGCACCGGGGCCAGCCCGAGCGAGTTTCGTGCCGGTTGTGAAGAAAAAGTGAATGATGTAGCCG
TCAAGTTGTCATAATAAGCTTTGCATCATCACCATCACCACGGATCCGCTGAAGCaCAAAATGATCCCCTGCTGCCG
GGATACTCGTTTAACGCCCATCTGGTGGCGGGTTTAACGCCGATTGAGGCCAACGGTTATCTCGATTTTTTTATCGA
CCGACCGCTGGGAATGAAAGGTTATATTCTCAATCTCACCATTCGCGGTCAGGGGGTGGTGAAAAATCAGGGACGAG
AATTTGTCTGCCGACCGGGTGATATTTTGCTGTTCCCGCCAGGAGAGATTCATCACTACGGTCGTCATCCGGAGGCT
CGCGAATGGTATCACCAGTGGGTTTACTTTCGTCCGCGCGCCTACTGGCATGAATGGCTTAACTGGCCGTCAATATT
TGCCAATACGGGTTTCTTTCGCCCGGATGAAGCGCACCAGCCGCATTTCAGCGACCTGTTTGGGCAAATCATTAACG
CCGGGCAAGGGGAAGGGCGCTATTCGGAGCTGCTGGCGATAAATCTGCTTGAGCAATTGTTACTGCGGCGCATGGAA
GCGATTAACGAGTCGCTCCATCCACCGATGGATAATCGGGTACGCGAGGCTTGTCAGTACATCAGCGATCACCTGGC
AGACAGCAATTTTGATATCGCCAGCGTCGCACAGCATGTTTGCTTGTCGCCGTCGCGTCTGTCACATCTTTTCCGCC
AGCAGTTAGGGATTAGCGTCTTAAGCTGGCGCGAGGACCAACGCATTAGTCAGGCGAAGCTGCTTTTGAGCACTACC
CGGATGCCTATCGCCACCGTCGGTCGCAATGTTGGTTTTGACGATCAACTCTATTTCTCGCGAGTATTTAAAAAATG
CACCGGGGCCAGCCCGAGCGAGTTTCGTGCCGGTTGTGAAGAAAAAGTGAATGATGTAGCCGTCAAGTTGTCATAAT
AAGCTTACaAGTaataCTGCAGAGAGAATATAAAAAGCCAGATTATTAATCCGGCTTTTTTATTATTTAGACGTCAG
GTGGCACTTTTCGGGGAAATGTGCGCGGAACCCCTATTTGTTTATTTTTCTAAATACATTCAAATATGTATCCGCTC
ATGAGACAATAACCCTGATAAATGCTTCAATAATATTGAAAAAGGAAGAGTATGAGTATTCAACATTTCCGTGTCGC
CCTTATTCCCTTTTTTGCGGCATTTTGCCTTCCTGTTTTTGCTCACCCAGAAACGCTGGTGAAAGTAAAAGATGCTG
AAGATCAGTTGGGTGCACGAGTGGGTTACATCGAACTGGATCTCAACAGCGGTAAGATCCTTGAGAGTTTTCGCCCC
GAAGAACGTTTTCCAATGATGAGCACTTTTAAAGTTCTGCTATGTGGCGCGGTATTATCCCGTATTGACGCCGGGCA
AGAGCAACTCGGTCGCCGCATACACTATTCTCAGAATGACTTGGTTGAGTACTCACCAGTCACAGAAAAGCATCTTA
CGGATGGCATGACAGTAAGAGAATTATGCAGTGCTGCCATAACCATGAGTGATAACACTGCGGCCAACTTACTTCTG
ACAACGATCGGAGGACCGAAGGAGCTAACCGCTTTTTTGCACAACATGGGGGATCATGTAACTCGCCTTGATCGTTG
GGAACCGGAGCTGAATGAAGCCATACCAAACGACGAGCGTGACACCACGATGCCTGTAGCAATGGCAACAACGTTGC
GCAAACTATTAACTGGCGAACTACTTACTCTAGCTTCCCGGCAACAATTAATAGACTGGATGGAGGCGGATAAAGTT
GCAGGACCACTTCTGCGCTCGGCCCTTCCGGCTGGCTGGTTTATTGCTGATAAATCTGGAGCCGGTGAGCGTGGtTC

TCGCGGTATCATTGCAGCACTGGGGCCAGATGGTAAGCCCTCCCGTATCGTAGTTATCTACACGACGGGGAGTCAGG
CAACTATGGATGAACGAAATAGACAGATCGCTGAGATAGGTGCCTCACTGATTAAGCATTGGTAAGTGACCAAACAG
GAAAAAACCGCCCTTAACATGGCCCGCTTTATCAGAAGCCAGACATTAACGCTTCTGGAGAAACTCAACGAGCTGGA
CGCGGATGAACAGGCAGACATCTGTGAATCGCTTCACGACCACGCTGATGAGCTTTACCGCAGCTGCCTCGCGCGTT
TCGGTGATGACGGTGAAAACCTCTGACTGTCAGACCAAGTTTACTCATATATACTTTAGATTGATTTAAAACTTCAT
TTTTAATTTAAAAGGATCTAGGTGAAGATCCTTTTTGATAATCTCATGACCAAAATCCCTTAACGTGAGTTTTCGTT
CCACTGAGCGTCAGACCCCGTAGAAAAGATCAAAGGATCTTCTTGAGATCCTTTTTTTCTGCGCGTAATCTGCTGCT
TGCAAACAAAAAAACCACCGCTACCAGCGGTGGTTTGTTTGCCGGATCAAGAGCTACCAACTCTTTTTCCGAAGGTA
ACTGGCTTCAGCAGAGCGCAGATACCAAATACTGTTCTTCTAGTGTAGCCGTAGTTAGGCCACCACTTCAAGAACTC
TGTAGCACCGCCTACATACCTCGCTCTGCTAATCCTGTTACCAGTGGCTGCTGCCAGTGGCGATAAGTCGTGTCTTA
CCGGGTTGGACTCAAGACGATAGTTACCGGATAAGGCGCAGCGGTCGGGCTGAACGGGGGGTTCGTGCACACAGCCC
AGCTTGGAGCGAACGACCTACACCGAACTGAGATACCTACAGCGTGAGCTATGAGAAAGCGCCACGCTTCCCGAAGG
GAGAAAGGCGGACAGGTATCCGGTAAGCGGCAGGGTCGGAACAGGAGAGCGCACGAGGGAGCTTCCAGGGGGAAACG
CCTGGTATCTTTATAGTCCTGTCGGGTTTCGCCACCTCTGACTTGAGCGTCGATTTTTGTGATGCTCGTCAGGGGGG
CGGAGCCTATGGAAA

**Figure S27. Plasmid map and nucleotide sequence of pAraC.**

AraC region is highlighted in yellow.

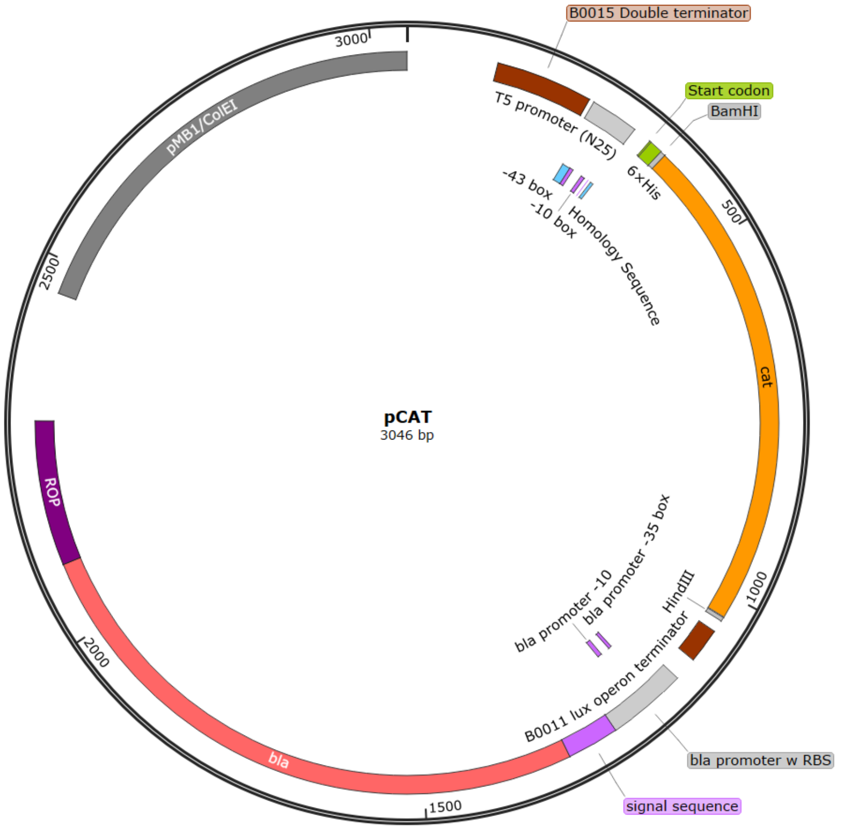


AACGCCAGCAACGCGGCCTTTTTACGGTTCCTGGCCTTTTGCTGGCCTTTTGCTCACATGTTCTTTCCTGCGTTATC
CCCTGATTCTGTGGATAACCGTATTACCGCCTTTGAGTGAGCTCCAGGCATCAAATAAAACGAAAGGCTCAGTCGAA
AGACTGGGCCTTTCGTTTTATCTGTTGTTTGTCGGTGAACGCTCTCTACTAGAGTCACACTGGCTCACCTTCGGGTG
GGCCTTTCTGCGTTTATAGGTACCAAGAATCATAAAAAATTTATTTGCTTTCAGGAAAATTTTTCTGTATAATAGAT
TCATAAATTTGAGGGCCCCCCTCCTTCAGGTCTAAAATGCAACAACAGTACTGCGATGAGTGGCAGGGCGGGGCGTA
ATAAGCTTACaAGTaataCTGCAGAGAGAATCATCACCATCACCACGGATCCGAGAAAAAAATCACTGGATATACCA
CCGTTGATATATCCCAATGGCATCGTAAAGAACATTTTGAGGCATTTCAGTCAGTTGCTCAATGTACCTATAACCAG
ACCGTTCAGCTGGATATTACGGCCTTTTTAAAGACCGTAAAGAAAAATAAGCACAAGTTTTATCCGGCCTTTATTCA
CATTCTTGCCCGCCTGATGAATGCTCATCCGGAATTCCGTATGGCAATGAAAGACGGTGAGCTGGTGATATGGGATA
GTGTTCACCCTTGTTACACCGTTTTCCATGAGCAAACTGAAACGTTTTCATCGCTCTGGAGTGAATACCACGACGAT
TTCCGGCAGTTTCTACACATATATTCGCAAGATGTGGCGTGTTACGGTGAAAACCTGGCCTATTTCCCTAAAGGGTT
TATTGAGAATATGTTTTTCGTCTCAGCCAATCCCTGGGTGAGTTTCACCAGTTTTGATTTAAACGTGGCCAATATGG
ACAACTTCTTCGCCCCCGTTTTCACCATGGGCAAATATTATACGCAAGGCGACAAGGTGCTGATGCCGCTGGCGATT
CAGGTTCATCATGCCGTCTGTGATGGCTTCCATGTCGGCAGAATGCTTAATGAATTACAACAGTACTGCGATGAGTG
GCAGGGCGGGGCGTAATAAGCTTACaAGTaataCTGCAGAGAGAATATAAAAAGCCAGATTATTAATCCGGCTTTTT
TATTATTTAGACGTCAGGTGGCACTTTTCGGGGAAATGTGCGCGGAACCCCTATTTGTTTATTTTTCTAAATACATT
CAAATATGTATCCGCTCATGAGACAATAACCCTGATAAATGCTTCAATAATATTGAAAAAGGAAGAGTATGAGTATT
CAACATTTCCGTGTCGCCCTTATTCCCTTTTTTGCGGCATTTTGCCTTCCTGTTTTTGCTCACCCAGAAACGCTGGT
GAAAGTAAAAGATGCTGAAGATCAGTTGGGTGCACGAGTGGGTTACATCGAACTGGATCTCAACAGCGGTAAGATCC
TTGAGAGTTTTCGCCCCGAAGAACGTTTTCCAATGATGAGCACTTTTAAAGTTCTGCTATGTGGCGCGGTATTATCC
CGTATTGACGCCGGGCAAGAGCAACTCGGTCGCCGCATACACTATTCTCAGAATGACTTGGTTGAGTACTCACCAGT
CACAGAAAAGCATCTTACGGATGGCATGACAGTAAGAGAATTATGCAGTGCTGCCATAACCATGAGTGATAACACTG
CGGCCAACTTACTTCTGACAACGATCGGAGGACCGAAGGAGCTAACCGCTTTTTTGCACAACATGGGGGATCATGTA
ACTCGCCTTGATCGTTGGGAACCGGAGCTGAATGAAGCCATACCAAACGACGAGCGTGACACCACGATGCCTGTAGC
AATGGCAACAACGTTGCGCAAACTATTAACTGGCGAACTACTTACTCTAGCTTCCCGGCAACAATTAATAGACTGGA
TGGAGGCGGATAAAGTTGCAGGACCACTTCTGCGCTCGGCCCTTCCGGCTGGCTGGTTTATTGCTGATAAATCTGGA
GCCGGTGAGCGTGGtTCTCGCGGTATCATTGCAGCACTGGGGCCAGATGGTAAGCCCTCCCGTATCGTAGTTATCTA
CACGACGGGGAGTCAGGCAACTATGGATGAACGAAATAGACAGATCGCTGAGATAGGTGCCTCACTGATTAAGCATT
GGTAAGTGACCAAACAGGAAAAAACCGCCCTTAACATGGCCCGCTTTATCAGAAGCCAGACATTAACGCTTCTGGAG

AAACTCAACGAGCTGGACGCGGATGAACAGGCAGACATCTGTGAATCGCTTCACGACCACGCTGATGAGCTTTACCG
CAGCTGCCTCGCGCGTTTCGGTGATGACGGTGAAAACCTCTGACTGTCAGACCAAGTTTACTCATATATACTTTAGA
TTGATTTAAAACTTCATTTTTAATTTAAAAGGATCTAGGTGAAGATCCTTTTTGATAATCTCATGACCAAAATCCCT
TAACGTGAGTTTTCGTTCCACTGAGCGTCAGACCCCGTAGAAAAGATCAAAGGATCTTCTTGAGATCCTTTTTTTCT
GCGCGTAATCTGCTGCTTGCAAACAAAAAAACCACCGCTACCAGCGGTGGTTTGTTTGCCGGATCAAGAGCTACCAA
CTCTTTTTCCGAAGGTAACTGGCTTCAGCAGAGCGCAGATACCAAATACTGTTCTTCTAGTGTAGCCGTAGTTAGGC
CACCACTTCAAGAACTCTGTAGCACCGCCTACATACCTCGCTCTGCTAATCCTGTTACCAGTGGCTGCTGCCAGTGG
CGATAAGTCGTGTCTTACCGGGTTGGACTCAAGACGATAGTTACCGGATAAGGCGCAGCGGTCGGGCTGAACGGGGG
GTTCGTGCACACAGCCCAGCTTGGAGCGAACGACCTACACCGAACTGAGATACCTACAGCGTGAGCTATGAGAAAGC
GCCACGCTTCCCGAAGGGAGAAAGGCGGACAGGTATCCGGTAAGCGGCAGGGTCGGAACAGGAGAGCGCACGAGGGA
GCTTCCAGGGGGAAACGCCTGGTATCTTTATAGTCCTGTCGGGTTTCGCCACCTCTGACTTGAGCGTCGATTTTTGT
GATGCTCGTCAGGGGGGCGGAGCCTATGGAAA

**Figure S28. Plasmid map and nucleotide sequence of pCAT.**

CAT region is highlighted in yellow.

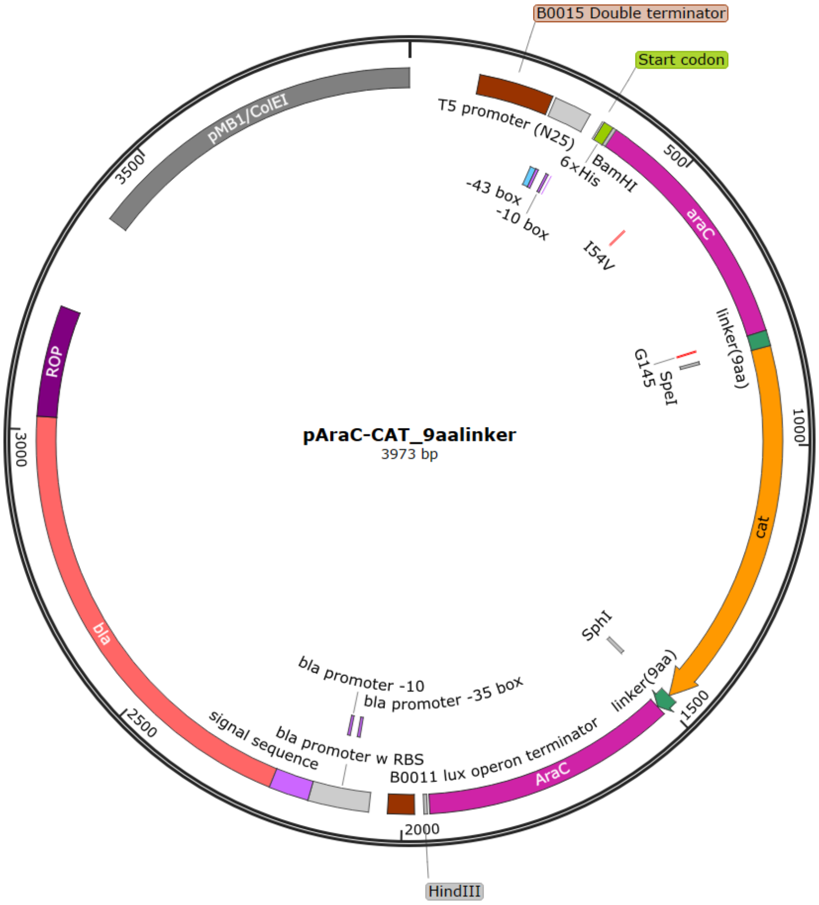


AACGCCAGCAACGCGGCCTTTTTACGGTTCCTGGCCTTTTGCTGGCCTTTTGCTCACATGTTCTTTCCTGCGTTATC
CCCTGATTCTGTGGATAACCGTATTACCGCCTTTGAGTGAGCTCCAGGCATCAAATAAAACGAAAGGCTCAGTCGAA
AGACTGGGCCTTTCGTTTTATCTGTTGTTTGTCGGTGAACGCTCTCTACTAGAGTCACACTGGCTCACCTTCGGGTG
GGCCTTTCTGCGTTTATAGGTACCAAGAATCATAAAAAATTTATTTGCTTTCAGGAAAATTTTTCTGTATAATAGAT
TCATAAATTTGAGGGCCCCCCTCCTTCAGGTCTAAAATGCATCATCACCATCACCACGGATCCGCTGAAGCaCAAAA
TGATCCCCTGCTGCCGGGATACTCGTTTAACGCCCATCTGGTGGCGGGTTTAACGCCGATTGAGGCCAACGGTTATC
TCGATTTTTTTATCGACCGACCGCTGGGAATGAAAGGTTATgTTCTCAATCTCACCATTCGCGGTCAGGGGGTGGTG
AAAAATCAGGGACGAGAATTTGTCTGCCGACCGGGTGATATTTTGCTGTTCCCGCCAGGAGAGATTCATCACTACGG
TCGTCATCCGGAGGCTCGCGAATGGTATCACCAGTGGGTTTACTTTCGTCCGCGCGCCTACTGGCATGAATGGCTTA
ACTGGCCGTCAATATTTGCCAATACGGGTTTCTTTCGCCCGGATGAAGCGCACCAGCCGCATTTCAGCGACCTGTTT
GGGCAAATCATTAACGCCGGGCAAGGGGAAGGGGGTGGcagTGGTGGTTCaGGcACTAGTGAGAAAAAAATCACTGG
ATATACCACCGTTGATATATCCCAATGGCATCGTAAAGAACATTTTGAGGCATTTCAGTCAGTTGCTCAATGTACCT
ATAACCAGACCGTTCAGCTGGATATTACGGCCTTTTTAAAGACCGTAAAGAAAAATAAGCACAAGTTTTATCCGGCC
TTTATTCACATTCTTGCCCGCCTGATGAATGCTCATCCGGAATTCCGTATGGCAATGAAAGACGGTGAGCTGGTGAT
ATGGGATAGTGTTCACCCTTGTTACACCGTTTTCCATGAGCAAACTGAAACGTTTTCATCGCTCTGGAGTGAATACC
ACGACGATTTCCGGCAGTTTCTACACATATATTCGCAAGATGTGGCGTGTTACGGTGAAAACCTGGCCTATTTCCCT
AAAGGGTTTATTGAGAATATGTTTTTCGTCTCAGCCAATCCCTGGGTGAGTTTCACCAGTTTTGATTTAAACGTGGC
CAATATGGACAACTTCTTCGCCCCCGTTTTCACCATGGGCAAATATTATACGCAAGGCGACAAGGTGCTGATGCCGC
TGGCGATTCAGGTTCATCATGCCGTtTGTGATGGCTTCCATGTCGGCAGAATGCTTAATGAATTACAACAGTACTGC
GATGAGTGGCAGGGCGGGGCGGCATGCggtagtggttctggcagtggcCGCTATTCGGAGCTGCTGGCGATAAATCT
GCTTGAGCAATTGTTACTGCGGCGCATGGAAGCGATTAACGAGTCGCTCCATCCACCGATGGATAATCGGGTACGCG
AGGCTTGTCAGTACATCAGCGATCACCTGGCAGACAGCAATTTTGATATCGCCAGCGTCGCACAGCATGTTTGCTTG
TCGCCGTCGCGTCTGTCACATCTTTTCCGCCAGCAGTTAGGGATTAGCGTCTTAAGCTGGCGCGAGGACCAACGCAT
TAGTCAGGCGAAGCTGCTTTTGAGCACTACCCGGATGCCTATCGCCACCGTCGGTCGCAATGTTGGTTTTGACGATC
AACTCTATTTCTCGCGAGTATTTAAAAAATGCACCGGGGCCAGCCCGAGCGAGTTTCGTGCCGGTTGTGAAGAAAAA
GTGAATGATGTAGCCGTCAAGTTGTCATAATAAGCTTACaAGTaataCTGCAGAGAGAATATAAAAAGCCAGATTAT
TAATCCGGCTTTTTTATTATTTAGACGTCAGGTGGCACTTTTCGGGGAAATGTGCGCGGAACCCCTATTTGTTTATT

TTTCTAAATACATTCAAATATGTATCCGCTCATGAGACAATAACCCTGATAAATGCTTCAATAATATTGAAAAAGGA
AGAGTATGAGTATTCAACATTTCCGTGTCGCCCTTATTCCCTTTTTTGCGGCATTTTGCCTTCCTGTTTTTGCTCAC
CCAGAAACGCTGGTGAAAGTAAAAGATGCTGAAGATCAGTTGGGTGCACGAGTGGGTTACATCGAACTGGATCTCAA
CAGCGGTAAGATCCTTGAGAGTTTTCGCCCCGAAGAACGTTTTCCAATGATGAGCACTTTTAAAGTTCTGCTATGTG
GCGCGGTATTATCCCGTATTGACGCCGGGCAAGAGCAACTCGGTCGCCGCATACACTATTCTCAGAATGACTTGGTT
GAGTACTCACCAGTCACAGAAAAGCATCTTACGGATGGCATGACAGTAAGAGAATTATGCAGTGCTGCCATAACCAT
GAGTGATAACACTGCGGCCAACTTACTTCTGACAACGATCGGAGGACCGAAGGAGCTAACCGCTTTTTTGCACAACA
TGGGGGATCATGTAACTCGCCTTGATCGTTGGGAACCGGAGCTGAATGAAGCCATACCAAACGACGAGCGTGACACC
ACGATGCCTGTAGCAATGGCAACAACGTTGCGCAAACTATTAACTGGCGAACTACTTACTCTAGCTTCCCGGCAACA
ATTAATAGACTGGATGGAGGCGGATAAAGTTGCAGGACCACTTCTGCGCTCGGCCCTTCCGGCTGGCTGGTTTATTG
CTGATAAATCTGGAGCCGGTGAGCGTGGtTCTCGCGGTATCATTGCAGCACTGGGGCCAGATGGTAAGCCCTCCCGT
ATCGTAGTTATCTACACGACGGGGAGTCAGGCAACTATGGATGAACGAAATAGACAGATCGCTGAGATAGGTGCCTC
ACTGATTAAGCATTGGTAAGTGACCAAACAGGAAAAAACCGCCCTTAACATGGCCCGCTTTATCAGAAGCCAGACAT
TAACGCTTCTGGAGAAACTCAACGAGCTGGACGCGGATGAACAGGCAGACATCTGTGAATCGCTTCACGACCACGCT
GATGAGCTTTACCGCAGCTGCCTCGCGCGTTTCGGTGATGACGGTGAAAACCTCTGACTGTCAGACCAAGTTTACTC
ATATATACTTTAGATTGATTTAAAACTTCATTTTTAATTTAAAAGGATCTAGGTGAAGATCCTTTTTGATAATCTCA
TGACCAAAATCCCTTAACGTGAGTTTTCGTTCCACTGAGCGTCAGACCCCGTAGAAAAGATCAAAGGATCTTCTTGA
GATCCTTTTTTTCTGCGCGTAATCTGCTGCTTGCAAACAAAAAAACCACCGCTACCAGCGGTGGTTTGTTTGCCGGA
TCAAGAGCTACCAACTCTTTTTCCGAAGGTAACTGGCTTCAGCAGAGCGCAGATACCAAATACTGTTCTTCTAGTGT
AGCCGTAGTTAGGCCACCACTTCAAGAACTCTGTAGCACCGCCTACATACCTCGCTCTGCTAATCCTGTTACCAGTG
GCTGCTGCCAGTGGCGATAAGTCGTGTCTTACCGGGTTGGACTCAAGACGATAGTTACCGGATAAGGCGCAGCGGTC
GGGCTGAACGGGGGGTTCGTGCACACAGCCCAGCTTGGAGCGAACGACCTACACCGAACTGAGATACCTACAGCGTG
AGCTATGAGAAAGCGCCACGCTTCCCGAAGGGAGAAAGGCGGACAGGTATCCGGTAAGCGGCAGGGTCGGAACAGGA
GAGCGCACGAGGGAGCTTCCAGGGGGAAACGCCTGGTATCTTTATAGTCCTGTCGGGTTTCGCCACCTCTGACTTGA
GCGTCGATTTTTGTGATGCTCGTCAGGGGGGCGGAGCCTATGGAAA

**Figure S29. Plasmid map and nucleotide sequence of pAraC-CAT_9aalinker.**

AraC$_{I54V}$-CAT region is highlighted in yellow.

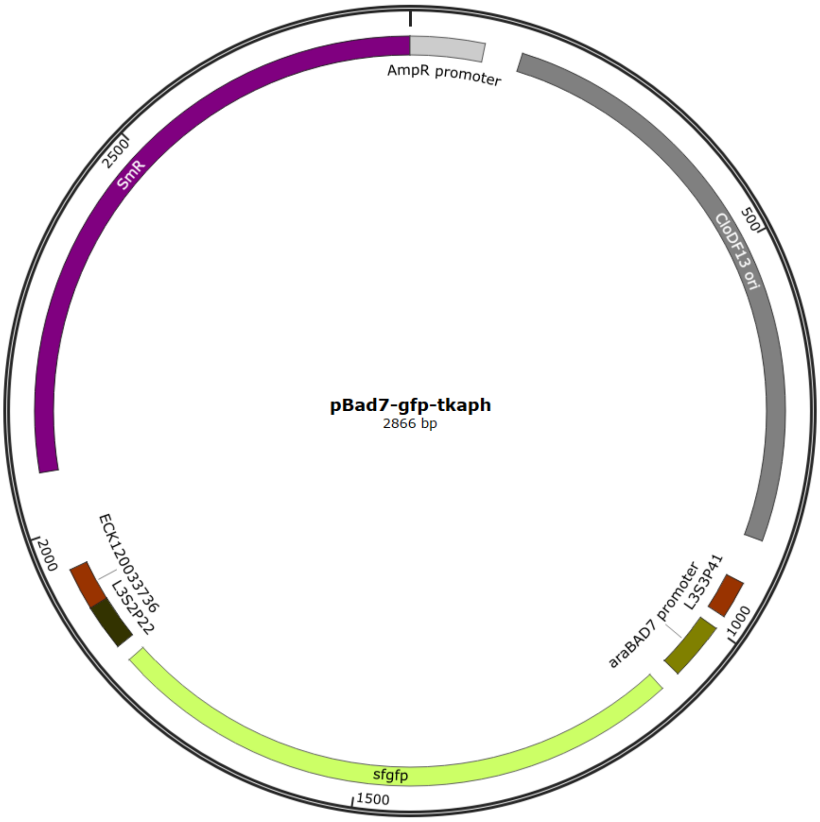


ACTCTTCCTTTTTCAATATTATTGAAGCATTTATCAGGGTTATTGTCTCATGAGCGGATACATATTTGAATGTATTT
AGAAAAATAAACAAATAGCTAGCTCACTCGGTCGCTACGCTCCGGGCGTGAGACTGCGGCGGGCGCTGCGGACACAT
ACAAAGTTACCCACAGATTCCGTGGATAAGCAGGGGACTAACATGTGAGGCAAAACAGCAGGGCCGCGCCGGTGGCG
TTTTTCCATAGGCTCCGCCCTCCTGCCAGAGTTCACATAAACAGACGCTTTTCCGGTGCATCTGTGGGAGCCGTGAG
GCTCAACCATGAATCTGACAGTACGGGCGAAACCCGACAGGACTTAAAGATCCCCACCGTTTCCGGCGGGTCGCTCC
CTCTTGCGCTCTCCTGTTCCGACCCTGCCGTTTACCGGATACCTGTTCCGCCTTTCTCCCTTACGGGAAGTGTGGCG
CTTTCTCATAGCTCACACACTGGTATCTCGGCTCGGTGTAGGTCGTTCGCTCCAAGCTGGGCTGTAAGCAAGAACTC
CCCGTTCAGCCCGACTGCTGCGCCTTATCCGGTAACTGTTCACTTGAGTCCAACCCGGAAAAGCACGGTAAAACGCC
ACTGGCAGCAGCCATTGGTAACTGGGAGTTCGCAGAGGATTTGTTTAGCTAAACACGCGGTTGCTCTTGAAGTGTGC
GCCAAAGTCCGGCTACACTGGAAGGACAGATTTGGTTGCTGTGCTCTGCGAAAGCCAGTTACCACGGTTAAGCAGTT
CCCCAACTGACTTAACCTTCGATCAAACCACCTCCCCAGGTGGTTTTTTCGTTTACAGGGCAAAAGATTACGCGCAG
AAAAAAAGGATCTCAAGAAGATCCTTTGATCTTTTCTACTGAACCGCTCTAGATTTCAGTGCAATTTATCTCTTCAA
ATGAGGATCCAAAAAAAAAAAAACACCCTAACGGGTGTTTTTTTTTTTTTGGTCTGCCTGACACCATGCAAGCTTTAG
CATTTTTATCCATAAGATTAGCATTTTTATCCATAGATCCTGGTACCGAATTCATTGTCTACTGTTTCTATAAATTT
GAGAGGGGAGCAACTAGTATGGGGTCTAAAGGCGAAGAACTGTTCACCGGCGTAGTTCCGATCCTGGTTGAACTGGA
CGGTGACGTTAATGGTCATAAGTTCTCTGTTCGTGGTGAAGGTGAGGGCGACGCGACCAACGGTAAACTGACCCTGA
AATTCATCTGCACCACTGGCAAACTGCCGGTTCCGTGGCCGACTCTGGTTACCACCCTGACCTATGGTGTTCAGTGC
TTCTCTCGTTACCCGGATCACATGAAACAGCACGACTTCTTCAAATCTGCGATGCCGGAGGGTTATGTTCAGGAACG
TACCATCTCTTTCAAGGATGACGGCACCTACAAAACCCGTGCCGAAGTTAAATTCGAGGGTGATACGCTGGTAAACC
GCATCGAACTGAAAGGTATCGACTTCAAAGAGGACGGTAATATCCTCGGTCACAAGCTGGAATACAACTTCAACTCT
CACAACGTTTACATCACCGCGGACAAACAGAAAAACGGTATCAAAGCGAACTTTAAGATCCGTCACAATGTTGAAGA
CGGCAGCGTTCAGCTCGCTGACCACTACCAACAAAATACCCCGATTGGCGACGGTCCGGTTCTGCTGCCGGACAACC
ACTATCTGTCTACCCAGTCTGTGCTCTCTAAGGACCCGAACGAGAAACGTGACCACATGGTGCTGCTGGAGTTCGTG
ACCGCAGCGGGCATCACGCACGGCATGGACGAACTGTACAAATGATAATAACATATGTCAGTCGACGAGCTCGGTAC
CAAATTCCAGAAAAGAGGCCGCGAAAGCGGCCTTTTTTCGTTTTGGTCCGCGCAATAAAAAAGCCCCCGGAAGGTGA
TCTTCCGGGGGCTTTCTCATGCGTTACAATATCGTTCACTCATCCAATCTCACTGACGGTTAGGCGTGCCTGGTACC
CTGCCCTGAACCGACGACCGGGTCATCGTGGCCGGATCTTGCGGCCCCTCGGCTTGAACGAATTGTTAGACATTATT
TGCCGACTACCTTGGTGATCTCGCCTTTCACGTAGTGGACAAATTCTTCCAACTGATCTGCGCGCGAGGCCAAGCGA
TCTTCTTCTTGTCCAAGATAAGCCTGTCTAGCTTCAAGTATGACGGGCTGATACTGGGCCGGCAGGCGCTCCATTGC
CCAGTCGGCAGCGACATCCTTCGGCGCGATTTTGCCGGTTACTGCGCTGTACCAAATGCGGGACAACGTAAGCACTA

CATTTCGCTCATCGCCAGCCCAGTCGGGCGGCGAGTTCCATAGCGTTAAGGTTTCATTTAGCGCCTCAAATAGATCC
TGTTCAGGAACCGGATCAAAGAGTTCCTCCGCCGCTGGACCTACCAAGGCAACGCTATGTTCTCTTGCTTTTGTCAG
CAAGATAGCCAGATCAATGTCGATCGTGGCTGGCTCGAAGATACCTGCAAGAATGTCATTGCGCTGCCATTCTCCAA
ATTGCAGTTCGCGCTTAGCTGGATAACGCCACGGAATGATGTCGTCGTGCACAACAATGGTGACTTCTACAGCGCGG
AGAATCTCGCTCTCTCCAGGGGAAGCCGAAGTTTCCAAAAGGTCGTTGATCAAAGCTCGCCGCGTTGTTTCATCAAG
CCTTACGGTCACCGTAACCAGCAAATCAATATCACTGTGTGGCTTCAGGCCGCCATCCACTGCGGAGCCGTACAAAT
GTACGGCCAGCAACGTCGGTTCGAGATGGCGCTCGATGACGCCAACTACCTCTGATAGTTGAGTCGATACTTCGGCG
ATCACCGCTTCCCTCAT

**Figure S30. Plasmid map and nucleotide sequence of pBAD7-gfp-tkaph**

$P_{BAD7}$ region is highlighted in green. sfGFPregion is highlighted in yellow.

**Table S1. The change in numbers of colonies with different Cm concentrations for each of the E. coli transformants of the libraries prepared in this study.**

| Library Name | Library size | [Cm] = 0 | [Cm] = 10 | [Cm] = 30 | [Cm] = 60 | [Cm] = 100 |
|---|---|---|---|---|---|---|
| (6 + 6) | $8.93\times10^4$ | ≒$10^4$ | 2178(21.8) | 756(7.56) | 423(4.23) | 113(1.13) |
| (3 + 3) | $1.34\times10^5$ | ≒$10^4$ | 1809(18.1) | 453(4.53) | 324(3.24) | 43(0.43) |
| (−4 + −4) | $3.12\times10^5$ | ≒$10^4$ | 2633(26.3) | 2100(2.10) | 1372(13.7) | 472(4.72) |
| (9 + 9)-G1 | $1.60\times10^5$ | ≒$10^4$ | 2109(21.1) | 634(6.34) | 312(3.12) | 102(1.02) |
| (9 + 9)-G2 | $2.16\times10^5$ | ≒$10^4$ | 1345(13.5) | 412(4.12) | 251(2.51) | 76(0.76) |
| (9 + 9)-G3 | $2.67\times10^5$ | ≒$10^4$ | 1023(10.2) | 213(2.13) | 142(1.42) | 12(0.12) |
| (9 + 9)-G4 | $3.87\times10^5$ | ≒$10^4$ | 219(2.19) | 143(1.43) | 54(0.54) | 0(0) |
| (9 + 9)-G5 | $2.02\times10^5$ | ≒$10^4$ | 78(0.78) | 23(0.23) | 4(0.04) | 0(0) |
| (9 + 9)-G5’ | $2.35\times10^5$ | ≒$10^4$ | 98(0.98) | 54(0.54) | 31(0.31) | 2(0) |
| (0 + 0)-G1 | $1.60\times10^5$ | ≒$10^4$ | >4000(>40) | >4000(>40) | 2032(20.3) | 540(5.40) |
| (0 + 0)-G2 | $4.12\times10^5$ | ≒$10^4$ | >4000(>40) | 2032(20.3) | 1423(14.2) | 439(4.39) |
| (0 + 0)-G3 | $3.12\times10^5$ | ≒$10^4$ | 987(9.87) | 802(8.02) | 491(4.91) | 354(3.54) |
| (0 + 0)-G4 | $9.12\times10^4$ | ≒$10^4$ | 423(4.23) | 257(2.57) | 52(0.52) | 32(0.32) |
| (0 + 0)-G5 | $7.89\times10^4$ | ≒$10^4$ | 180(1.80) | 75(0.75) | 43(0.43) | 0(0) |
| (0 + 0)-G5’ | $6.29\times10^4$ | ≒$10^4$ | 94(0.94) | 65(0.65) | 26(0.26) | 0(0) |

**Table S2. ΔΔG calculations for eight high-FD score variants (9 + 9)-G1$^{H}$.**

| FD-Score | N-side | C-side | Total [kJ/mol] |
|---|---|---|---|
| 0.4077 | Met-Asn | Cys-Gly | 4.366 |
| 0.3176 | Met-Asn | Cys-Val | 4.459 |
| 0.2998 | Glu-Ser | Arg-Gln | 8.198 |
| 0.2860 | Ser-His | Lys-Val | 8.749 |
| 0.2841 | Met-Asn | Cys-Lys | 8.368 |
| 0.2798 | Gly-Thr | Leu-Asn | 7.982 |
| 0.2741 | Gln-Gln | Ala-Val | 5.514 |
| 0.2737 | Ser-Gln | Arg-Ser | 4.673 |

**Table S3. ΔΔG calculations for eight mutants with low FD-scores in the 9aa linker library.**

| FD-Score | N-side | C-side | Total [kJ/mol] |
|---|---|---|---|
| －0.04643 | Gly-His | Asp-Phe | −0.8174 |
| −0.04502 | Thr−Leu | His−Ala | −0.6876 |
| −0.02385 | Tyr−Thr | Arg−Ala | 0.1093 |
| −0.002069 | Arg−Asp | Gly−Thr | 0.4223 |
| −0.0004076 | Thr−His | Lys−Glu | −0.3248 |
| 0.003186 | Val−Ile | Ala−Gly | 0.3355 |
| 0.005050 | Val−Asn | Gly−His | 0.1908 |
| 0.01172 | Pro−Gly | Tyr−Gly | 0.19364 |

**Table S4. ΔΔG calculations for eight mutants with the low FD-scores in the (0+0) linker library.**

| FD−Score | N-side | C-side | Total [kJ/mol] |
|---|---|---|---|
| −0.3781 | Cys−Lys | Val−Asn | 12.65 |
| −0.3642 | Pro−Asn | Ser−Val | 9.952 |
| −0.3592 | Cys−Ser | Val−Ala | 8.957 |
| −0.3401 | Cys−Lys | Ala−Cys | 7.613 |
| −0.3217 | Pro−Lys | Cys−Val | 11.11 |
| −0.2797 | Ser−His | Val−His | 18.15 |
| −0.2772 | Ala−Lys | Ser−Val | 12.56 |
| −0.2386 | Ala−Asn | Val−Cys | 11.14 |

**Table S5. ΔΔG calculations for eight mutants with the high FD-scores in the (0+0) linker library.**

| FD−Score | N-side | C-side | Total [kJ/mol] |
|---|---|---|---|
| 0.01610 | Gly−His | Ile−Thr | 4.357 |
| 0 | Leu−Gly | Tyr−Leu | 2.726 |
| −0.01801 | Arg−Arg | Val−Gln | 2.693 |
| −0.01832 | Ala−Tyr | Gly−Gly | 4.955 |
| −0.02635 | Gly−Glu | Phe−Ala | 4.273 |
| −0.02852 | Leu−Asn | Lys−Ala | 3.055 |
| −0.03043 | Ser−Arg | Ala−Asn | 2.007 |
| −0.03560 | Val−Ala | Met−His | 0.5170 |

**Table S6. Escherichia coli strains used in this study.**

| Name | Genotype | Source |
|---|---|---|
| **JW0063** | F⁻, *Δ(araC-araB)567, ΔaraC771::kan, ΔlacZ4787*(::rrnB-3), *λ⁻, rph-1, Δ(rhaD-rhaB)568, hsdR514* | Ref. 1 |
| **JW0063(ΔaraC)** | F⁻, *Δ(araC-araB)567, ΔaraC, ΔlacZ4787(*::rrnB-3), *λ⁻, rph-1, Δ(rhaD-rhaB)568, hsdR514* | This study |
| **XL10-Gold** | Tet$^r$, *ΔmcrA183, Δ(mcrCB-hsdSMR-mrr)173, endA1, supE44, thi-1, recA1, gyrA96, relA1, lac,* Hte [F$^l$, *proAB, lacI$^q$ZΔM15,* Tn10(Tet$^r$), Tn5(Kan$^r$), Amy] | Commercial strain |

**Table S7. Plasmids used in this study**

| Name | Based on vector | Origin | Marker | Source |
|---|---|---|---|---|
| pAraC | pHRA | ColE1 | $Amp^R$ | Laboratory stock |
| pCAT | pHRA | ColE1 | $Amp^R$ | Laboratory stock |
| pAraC−CAT_9aalinker | pHRA | ColE1 | $Amp^R$ | This study |
| pBad7−gfp−tkaph | pCloDF | CloDF13 | $Strep^R$ | This study |

**Table S8. Primers used in this study**

| Name | Sequence（5’ to 3’） |
| --- | --- |
| Ins Fwd | ATTACCGCCTTTGAGTGAGC |
| Ins Rev | GATAATACCGCGCCACATAGC |
| Vec Fwd | AGCGTCGCACAGCATGTTTG |
| Vec Rev | TGACCGCGAATGGTGAGATTG |
| 9aa N−Fwd | TTTTGGTCTCtGAGAAAAAAATCACTGGATATACCACCG |
| 9aa N Rev | TTTTGGTCTCtTCTCMNNMNNgCCtGAACCACCActgC |
| 6aa N Rev | TTTTGGTCTCtCTCTMNNMNNACCActgCCACCCCCTTC |
| 3aa N Rev | TTTTGGTCTCtTCTCMNNMNNACCCCCTTCCCCTTGCC |
| 0aa N Rev | TTTTGGTCTCtTCTCMNNMNNCCCTTGCCCGGCGTTAATG |
| −4aa N Fwd | TTTTGGTCTCtAGGGNNKNNKTATACCACCGTTGATATATC CCAATGG |
| −6aa N Fwd | TTTTGGTCTCtAGGGNNKNNKACCGTTGATATATCCCAATG GCATC |
| −4aa N Rev | TTTTGGTCTCtCCCTTGCCCGGCGTTAATG |
| 9aa C Fwd | TTTTGGTCTCtGGCGNNKNNKggtagtggttctggcagtg |
| 6aa C Fwd | TTTTGGTCTCtGGCGNNKNNKtctggcagtggcCGCTATTC |
| 3aa C Fwd | TTTTGGTCTCtGGCGNNKNNKggcCGCTATTCGGAGC |
| 0aa C Fwd | TTTTGGTCTCtGGCGNNKNNKTCGGAGCTGCTGGCGATAA ATC |
| 9aa C Rev | TTTTGGTCTCtCGCCCCGCCCTGC |
| −4aa C Fwd | TTTTGGTCTCtTCGGAGCTGCTGGCGATAAATC |
| −4aa C Rev | TTTTGGTCTCtGGTTMNNMNNATCGCAGTACTGTTGTAAT TCATTAAGCATTC |
| −6aa C Rev | TTTTGGTCTCtGGTTMNNMNNGTACTGTTGTAATTCATTA AGCATTCTGCC |
